# Flavoproteins as native and genetically encoded spin probes for *in cell* ESR spectroscopy

**DOI:** 10.1101/2024.12.18.628722

**Authors:** Timothée Chauviré, Siddarth Chandrasekaran, Robert Dunleavy, Jack H. Freed, Brian R. Crane

## Abstract

Flavin cofactors are attractive Electron Spin Resonance (ESR) probes for proteins because cellular reductants and light can generate their semiquinone states. We have used ESR spectroscopy to study the bacterial transmembrane aerotaxis receptor (Aer) in its native *Escherichia coli* membrane environment. Optimization of the spectroscopic (electronic relaxation times) and cell growth (isotopic labeling) conditions allowed for measurements of Aer with its partners - the histidine kinase (CheA) and the coupling protein (CheW) - in native signaling arrays. Continuous-wave ESR measurements at room temperature showed a rigid Aer flavin immobilized in the cofactor pocket and Q-band electron nuclear double resonance (ENDOR) measurements identified a predominant anionic semiquinone radical state *in cell*. Q-band four-pulse double electron-electron resonance (4P-DEER) measurements indicated a 4.1 nm distance between the two flavins of an Aer homodimer, consistent with previous *in vitro* measurements, but also revealed additional separations *in cell* indicative of chemoreceptor arrays, not previously observed for Aer. For general application, we further developed a genetically encoded Light-Oxygen and Voltage (LOV) domain for incorporation into target proteins as an ESR probe of structural properties *in cell*. This approach provides a framework to elucidate protein oligomeric states and conformations that are difficult to reproduce *in vitro*.

## INTRODUCTION

Flavins constitute redox-active protein cofactors that participate in a wide range of functions including catalysis^1^, light sensing^2^, and signal transduction^3^. In fact, flavoprotein genes constitute between 1 and 3 percent of both prokaryote and eukaryote genomes^4,5^. The ability to undergo one- and two-proton-coupled electron transfer reactions, provides versatility in enzymatic catalysis and allows for a diverse range of functions including oxidoreductase, transferase, and lyase activities^6^. Flavoproteins also comprise major classes of blue-light photoreceptors in the form of cryptochromes, Light Oxygen Voltage (LOV) domains and Blue-Light using FAD (BLUF) domains, which sense and respond to environmental input in many different organisms^7^. Similarly, flavoproteins play a major role in signaling pathways, particularly in bacterial systems, where flavin bound LOV and Per-Arnt-Sim (PAS) domains regulate kinase activity, DNA binding, and enzymatic activity^8^. This diversity of reactivity makes flavoproteins indispensable for a broad range of physiological functions, including oxidative damage response^9^, small-molecule metabolism^1^, and circadian rhythms^10^.

The term flavin is used to describe compounds containing 7,8-dimethyl-10-alkylisoalloxazine, which are characteristically yellow in color. Flavin cofactors are usually found either as the flavin adenine dinucleotide (FAD) or the flavin mononucleotide (FMN), which are both metabolites of riboflavin (Vitamin B2) and differ by the R group attached to the N(10) atom^6^. The majority of flavoproteins (75 %) utilize FAD as a cofactor, whereas the remaining (25 %) utilize FMN^6^. Only a few proteins directly use riboflavin as a cofactor; for example, the sodium pumping NADH:ubiquinone oxidoreductases of certain pathogenic bacteria, such as *Vibrio cholerae*^11^.

Flavins assume three different redox states, the quinone (oxidized), semiquinone (one-electron reduction), and hydroquinone (two-electron reduction). Depending on the cofactor environment, the semiquinone state is encountered in two different protonation states: the neutral or the anionic semiquinone (NSQ or ASQ, respectively). In solution, the semiquinone state of free flavin is unstable and is rapidly reduced to the hydroquinone^12^. However, flavoproteins stabilize either the neutral or anionic semiquinone state, enabling both photochemical and catalytic properties distinct from the free cofactor^13^.

The semiquinone state with its unpaired electron allows study by electron spin resonance (ESR) spectroscopy, which has been accomplished on both natural and synthetic semiquinones^13,14^. Because of their instability in aqueous solution, flavin semiquinone radicals have mostly been studied as protein-bound cofactors^13^, although an agarose gel matrix has been used to stabilize the FMN semiquinone state, allowing stable radicals for days under aerobic conditions^15^. Protein-bound flavin semiquinones ranging from glucose oxidase^16^ to light sensing cryptochromes^17^ have been analyzed using a combination of continuous wave ESR (cw-ESR) and pulsed ESR spectroscopy: pulsed dipolar spectroscopy (PDS), electron nuclear double resonance (ENDOR) or electron spin echo envelope modulation (ESEEM). Flavin semiquinones display characteristically broad features at X-band cw-ESR, with a linewidth of 1.5 mT for ASQs and 1.8-2.0 mT for NSQ due to inhomogeneous broadening from nitrogen and proton hyperfine interactions. Higher frequencies have been employed to accurately resolve g-factor tensor values using cw-ESR^16^. Pulsed ENDOR methods such as Davies ENDOR are particularly valuable to measure the large hyperfine couplings of flavin semiquinones^18^. Additionally, three and four-pulse ESEEM methods have been used to characterize hyperfine coupling constants (hfccs) in select flavoproteins^19^.

We sought to investigate the use of flavin semiquinone radicals for *in cell* measurement with ESR spectroscopy. Flavin cofactors offer several advantages for this application as they are tightly bound in their cofactor pocket, with dissociation constants in the nanomolar range^20^. Flavin binding domains are also small enough to be attached to the termini of a protein of interest; for example, FMN-binding LOV domains average only 110 residues^21^. Because FAD and FMN are natural metabolic cofactors, addition of exogenous cofactor is unnecessary for protein expression. The formation of ASQ or NSQ radicals can be induced by either chemical reductants or through illumination with blue light (λ = 420-480 nm), as some flavoproteins readily undergo flavin photoreduction. For *in cell* measurements, reduction can be accomplished in the reducing environment of the cell for certain proteins and enhanced by light.

Aer is an FAD-containing transmembrane protein in *E. coli* responsible for movement towards environments rich in terminal acceptors of the electron-transport chain, such as oxygen under aerobic conditions (i.e. aerotaxis)^22^. Aer is an obligate homodimer with multiple domains, an N-terminal FAD-binding PAS domain, a two-helix transmembrane region, a HAMP signaling domain (<u>h</u>istidine kinases, <u>a</u>denylyl cyclases, <u>m</u>ethyl-accepting chemotaxis proteins, <u>p</u>hosphatases), and a C-terminal kinase control domain^22^ (**Figure 1a, b**). Aer indirectly senses oxygen by measuring the redox environment of the cell through reactivity of its FAD-bound cofactor^23–26^. Changes in the FAD redox state induce conformational changes to the cytoplasmic kinase control domain, which in turn regulates activity of the histidine kinase CheA^27^. CheA phosphorylates a response regulator (CheY), which binds to the flagellar motor and switches the sense of motor rotation. Oxidized FAD in Aer supports CheA activation, whereas reduction to the anionic semiquinone inhibits CheA^28^. Physiological experiments suggest a third, hydroquinone form of FAD also producing CheA activation^23^, but the reduced hydroquinone state is not stable in purified Aer under aerobic conditions^28^. Within cells, *E. coli* chemoreceptors that regulate CheA associate as trimers-of-dimers (TODs) and extended molecular arrays of hexagonal symmetry in the membrane^29–33^. The cytoplasmic HAMP and kinase-control regions of Aer are closely related to those of chemoreceptors, and Aer does appear to form higher order oligomers *in vitro*^27^; however, it is unclear if Aer also forms extended arrays in cells.

**Figure 1:**
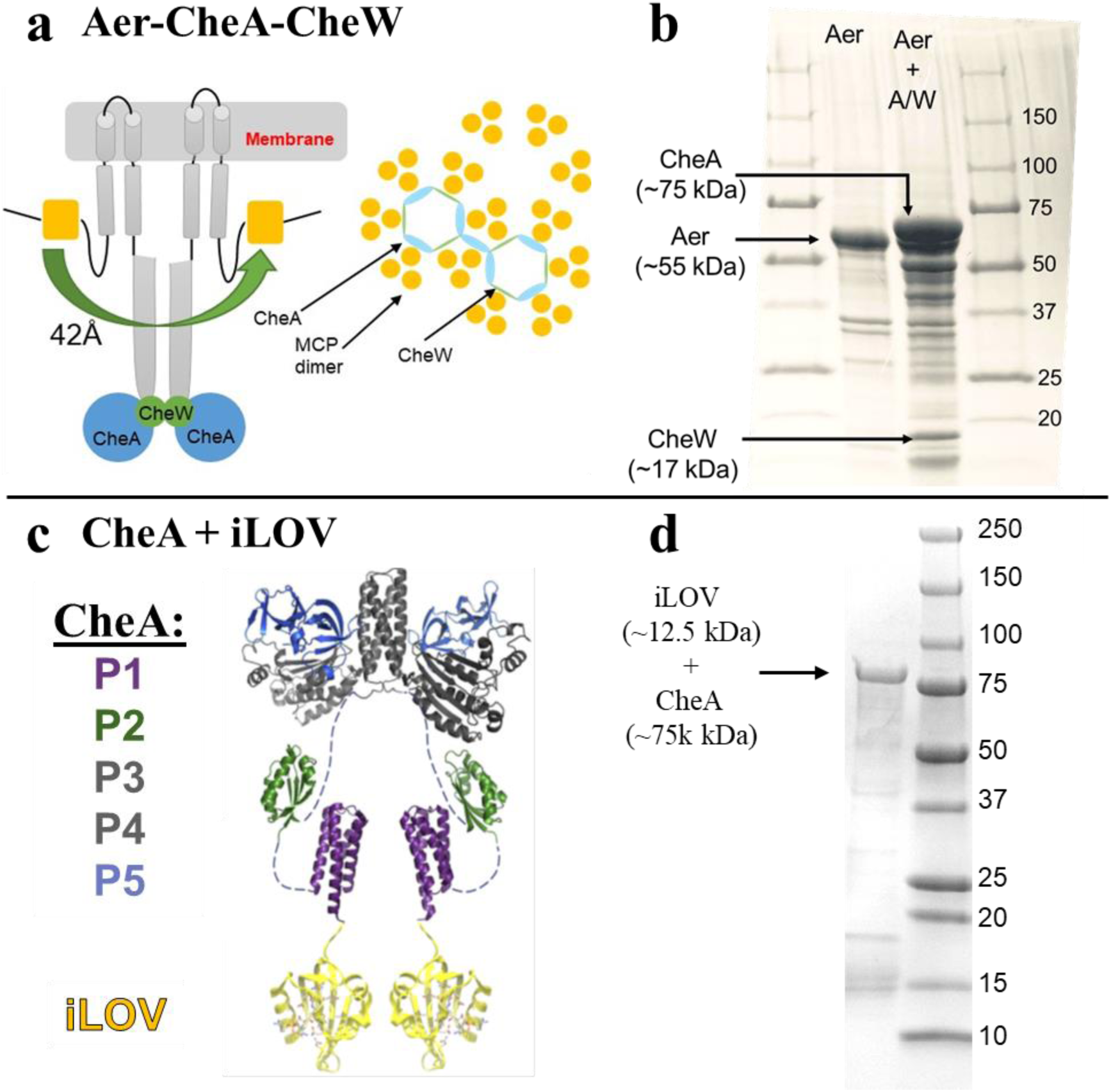
The two different chemosensory systems analyzed by ESR: Aer with an endogenous flavin center and CheA with an extraneous flavin center supplied by a small flavoprotein iLOV. (a) Representation of Aer and its binding partners, CheA & CheW, on the inner membrane of *E. coli* as viewed parallel to the membrane (left) and as viewed normal to the membrane (right); CheW and the P5 domain of CheA form hexameric rings that bind receptor trimers-of-dimers (TODs) (b) SDS-PAGE gel of the membrane fraction of BL21-DE3 cells overexpressing Aer (only) or Aer with CheA/CheW. (c) Representation of the fusion of the FMN-binding iLOV (∼12.5kDa) to the N-terminus of CheA (P1 Domain); the P1 domains of CheA are predicted to “lock-down” and dimerize when incorporated into receptor arrays. (d) SDS-PAGE gel of the membrane fraction of BL21-DE3 cells overexpressing CheA-iLOV.

We have previously used light-generated semiquinone radicals to determine the positioning of the Aer PAS sensing domains when full-length Aer was incorporated into nanodiscs^27^. Herein, we investigate the Aer receptor *in cell* both to provide native information about Aer and to elucidate general parameters for *in cell* flavoprotein ESR analysis. Previously, *in cell* ESR on flavoproteins performed for Arabidopsis thaliana cryptochrome 1 and 2 (cw-ESR, transient ESR and ENDOR)^18,34–38^ and Drosophila melanogaster cryptochrome (X-band cw-ESR, ENDOR)^39^, both in insect cells (Sf21) provided direct evidence of ASQ and NSQ formation, respectively, upon illumination with blue light^18,34–39^. In addition to cw-ESR and ENDOR, pulsed dipolar spectroscopy (PDS), such as four-pulse double electron-electron resonance (4P-DEER), has the potential to directly report on native protein conformations within cells. This approach could be particularly advantageous if the environment and interaction partners of the protein of interest are unknown and/or difficult to reconstitute *in vitro*.

*In cell* PDS measurements are challenging for several reasons: 1) delivery of spin-labeled biomolecules into cells is generally difficult, 2) spin labels are usually unstable in the reducing environment of the cell, 3) the spin-spin relaxation time (T_2_), also known as the phase memory time (T_m_) of the spin label, is too short due to various relaxation pathways present in the crowded cellular environment. To date, a few types of paramagnetic species have been employed in PDS measurements *in cell*, these include nitroxide spin labels^40–47^, gadolinium complexes^48–53^, trityl derivatives^54–57^, tyrosine radicals^58^ and copper-NTA complexes^59,60^; however, to our knowledge, flavins have yet to be used as paramagnetic centers for *in cell* PDS measurements. In this study, we show that both native flavoproteins and genetically engineered flavoprotein domains can be successfully used for *in cell* distance measurements. We investigate two systems involved in the *E. coli* chemotaxis signaling pathway: 1) the energy sensor Aer co-expressed with the scaffold protein CheW and the histidine kinase domain CheA (**Figure 1a,b**), 2) the histidine kinase CheA fused with a small (12.5 kDa) engineered Light Oxygen Voltage (LOV) domain (**Figure 1c,d**). Whole cell isotope labeling with ^15^N and ^2^H and temperature dependent studies were employed in order to optimize relaxation properties for 4P-DEER measurements. Overall, these findings provide a template for further *in cell* studies of both native flavoproteins and flavoprotein-based probes.

## RESULTS AND DISCUSSION

### Continuous Wave ESR and pulsed ENDOR of Aer *in cell*

The structural changes to Aer responsible for its energy taxis behavior are not well understood, particularly because it is difficult to isolate and characterize Aer complexes *in vitro*. Isolation of Aer in detergents, nanodiscs and lipodiscs^27,28^ yielded Aer in primarily a dimeric state, with only minor evidence for the TODs responsible for cooperative signaling in *E. coli*^61^. All of the experiments presented here used *E. coli* BL21 (DE3) cells as the host bacterial system. The *aer* gene was expressed from the T7 promoter of pET-28a plasmid, either alone or with its signaling partners CheA/CheW (**Figure 1b, Figure S1)**. After growth, a cell pellet was immediately harvested, inserted into an ESR capillary and studied by cw-ESR at room temperature to determine the concentration of semiquinone radical. The capillary was then flash-frozen to carry out pulsed ESR measurements.

Cw-ESR of over-expressed Aer in intact *E. coli* cells produced a stable signal of a characteristic semiquinone radical (**Figure 2a**), whereas the similar expression of the non-FAD binding aspartate receptor Tar revealed no such signal. The cytosol of *E. coli* is a reducing environment^62–64^ with a redox potential of ∼ -260 mV. The Aer redox potential in reconstituted systems is slightly lower at -290 mV^28^. Given that cellular redox state depends on cellular environment, metabolic conditions and is not generally at equilibrium^65^, it was unclear what the majority redox state of the Aer flavin would be in the cell. These measurements revealed that the anionic semiquinone state predominates.

**Figure 2:**
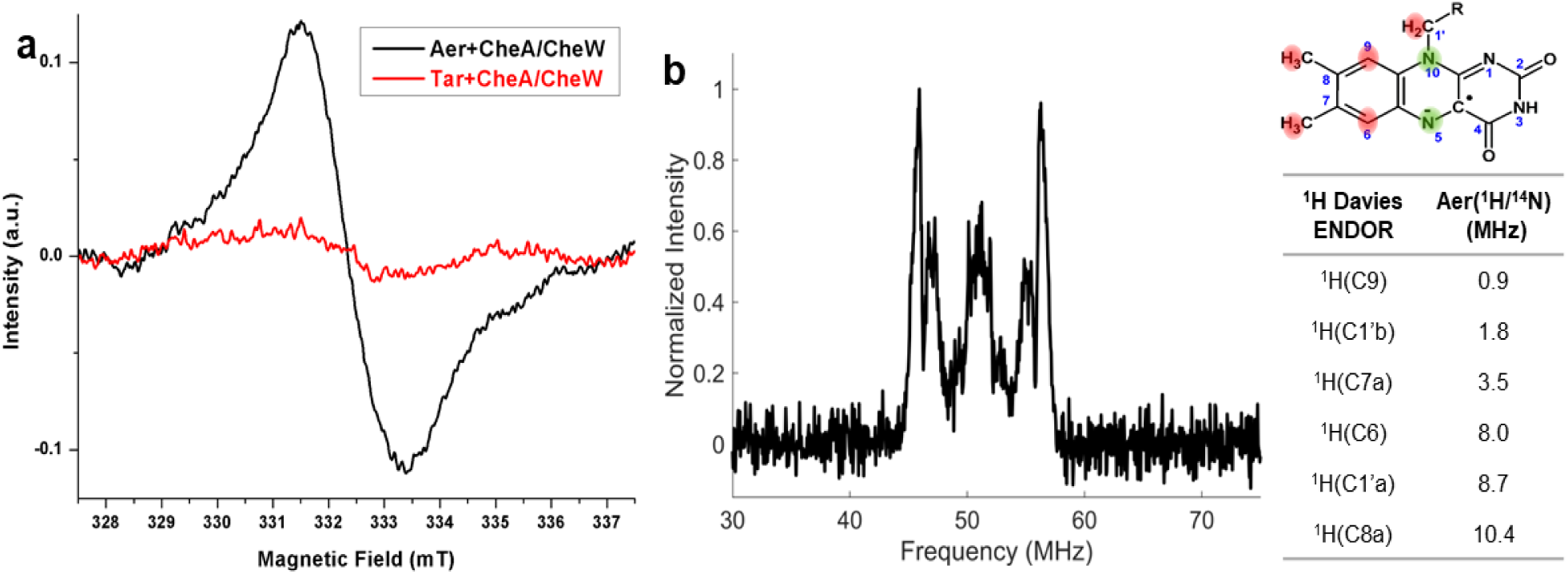
X-band continuous wave ESR and Q-Band ENDOR spectra of Aer *in cell*. (a) X-band cw-ESR spectra of BL21 (DE3) cells overexpressing CheA/CheW along with Aer (black curve) or Tar (red curve); (b) Q-band ^1^H Davies ENDOR spectra of Aer co-expressed with CheA/CheW in BL21 (DE3) cells observed at T = 150 K; The structure of the flavin isoalloxazine ring with the flavin protons highlighted in red along with their corresponding ENDOR frequencies below.

Proteomics studies have shown that *E. coli* BL21 (DE3) cells do not express the major protein components of chemotaxis (CheA, CheW, Tsr, Tar); however low levels of the receptors Trg and Aer were detected^66^. Aer and Trg are minor chemoreceptors in *E. coli*, accounting for less than 3-5 % of the total amount of chemoreceptors in a motile *E. coli* cell^67^. Accordingly *E. coli* BL21 (DE3) cells grown without plasmid or with a plasmid containing CheA and CheW gave a residual cw-ESR signal of ∼3 μM spin, which could be attributed to native Aer or other flavoproteins (**Figure S1**). Aer overexpression increased the concentration of semiquinone radical to 16 μM, whereas Aer overexpression with CheA and CheW produced a further increase to 26 μM. Earlier studies indicate that Aer expression leads to a concomitant overexpression of FAD^68,69^, raising the possibility that a portion of the signal-producing FAD may have been unbound to Aer. However, FAD semiquinones (both ASQ and NSQ) rapidly decay in solution^70^ unless stabilized by a protein matrix. Furthermore, experiments carried out on Aer variant that do not bind flavin^68,71,72^ (Aer Tyr93His/Cys193His/Cys203His) showed a substantial reduction of the *in cell* semiquinone signal (**Figure S2**). Thus, the observed semiquinone signal by cw-ESR owes primarily to FAD bound by the Aer receptor.

To further probe the radical state of Aer *in cell*, we performed Q-band ^1^H Davies ENDOR measurements (**Figure 2b**) at T = 150 K. The primary hfcc of 10.4 MHz corresponds to the H(8α) proton, whereas the hfccs observed at 8.7 and 8.0 MHz corresponds to the H(1’) and H(6) protons, respectively. The relatively high value of H(8α)^18^ and the absence of the large hfcc (∼25 MHz) corresponding to H(5) confirmed that Aer forms an ASQ radical *in cell*, as has been observed previously in preliminary overexpression experiments^28^. These values agree well with those from other ASQ-forming flavoproteins, including *Drosophila melanogaster* cryptochrome^17,39^ and *Aspergillus niger* glucose oxidase^16^.

### Four-pulse DEER of Aer *in cell*

As Aer is presumed to form a homodimer in the cytoplasmic membrane^23,27^, a 4.2 nm inter-subunit distance between flavin semiquinone radicals should be observed by 4P-DEER *in cell*. Indeed, 4P-DEER measurements for *in cell* Aer expressed alone, co-expressed with CheA/CheW, and *in vitro* Aer purified and solubilized in detergent (**Table S1**) showed a similar distance at 4.1 nm (**Figure 3a, b**). Thus, the cellular environment does not perturb the PAS domain positioning of the homodimer, and neither does association with CheA and CheW. The distance distribution for the radical pair is narrow with a width ranging from 0.2 to 0.3 nm (Full width half maximum (FWHM)), which indicates that the flavin is rigidly bound in the Aer PAS domain pocket on the nanosecond timescale of the experiment (**Figure 3b**). Interestingly, additional distances (3.3 nm and 4.9 nm) were observed for Aer-CheA-CheW *in cell*. Whereas the 4.9 nm distance should be interpreted with caution (as the background subtraction used can interfere and lead to artefactual signals at longer distances^73–75^), the shorter distance indicated association of Aer dimers in the cell membrane. Such a short distance is unlikely to arise from a single dimer because the HAMP domains that separate the PAS domains prevent a flavin-to-flavin separation closer than 4.1 nm. Interestingly, the shorter distance was absent when Aer was expressed without CheA/CheW, thereby indicating that the signaling partners stabilize assembly of Aer dimers into higher order structures (**Figure S3**).

**Figure 3:**
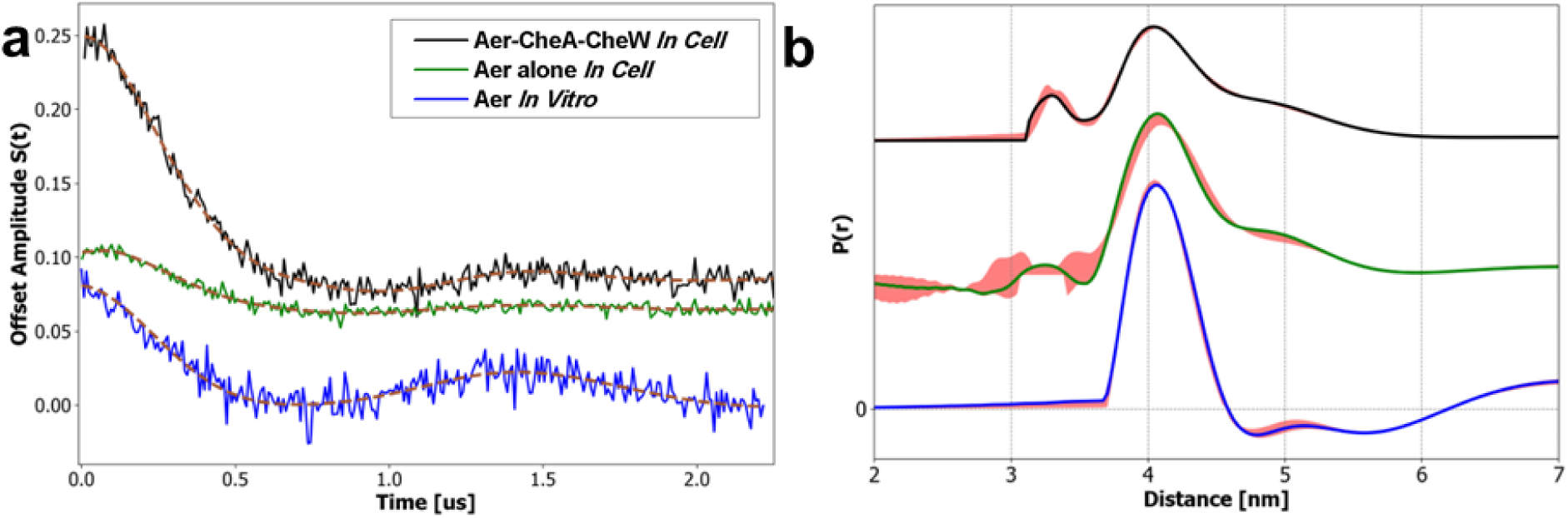
4P-DEER spectra of *in cell* Aer at T = 150 K. (a) Time domain and (b) distance domain distribution between anionic semiquinone radicals of Aer in *E. coli* BL21 (DE3) cells co-expressed with CheA/CheW (black upper curve), of Aer alone in *E. coli* BL21 (DE3) cells (green middle curve), and of purified Aer solubilized in detergent (blue lower curve). The distance domain was obtained by using the SF-SVD method^76^ and the time domain reconstructed spectrum is shown by a brown dashed line. Errors in the distance distributions represented by red shading and calculated as described in Srivastava et al.^72^.

The time domain 4P-DEER signal for Aer *in cell* when co-expressed with CheA/CheW (**Figure 3a**) displayed a modulation depth of 15.7 % (**Table S1**), which is typical for flavin anionic semiquinone radicals at Q-band^77^. Overexpression of Aer in the absence of CheA/CheW, caused a significant drop in the modulation depth (to 3.6 %) (**Table S1**). We carried out multiple measurements to confirm that this drop in modulation depth was significant (**Table S2**). Overexpression of Aer without its binding partners (CheA/CheW) likely leads to protein aggregation and therefore incomplete incorporation into the inner membrane of *E. coli.* Indeed, when expressed with CheA/CheW^68^, Aer was associated primarily in the membrane fraction (high-speed centrifugation fraction), but in the absence of CheA/CheW Aer was also associated with inclusion bodies and cellular debris (low-speed centrifugation fraction)^27^.

### Isotopic labeling of Aer *in cell*

For any spin system, an understanding of the relaxation properties (T_1_ and T_m_) is essential for optimal ESR data acquisition. For example, the longest distance that can be measured by 4P-DEER is T_m_ limited^75^. Phase memory times (T_m_) of flavin cofactors bound to proteins are relatively short (around 2 μs), and given that the dipolar oscillation frequency is inversely proportional to the cube of the distance between the spin labels, the maximum distances that can be measured at T_m_ ∼ 2 μs are limited to ∼ 5-6 nm^75^. Nonetheless, macromolecular complexes formed *in cell* can be separated by much larger distances (∼ 10’s of nm). Hence, lengthening the phase memory time of these flavin radicals will allow for measurements at longer evolution times.

Deuteration in the vicinity of the spin center substantially lower the rate of spin relaxation. For the case of a nitroxide radical, deuterated buffer can increase spin relaxation times by a factor of ∼ 2-3^78,79^, whereas a fully deuterated protein can improve the relaxation properties of the radical by a factor of ∼ 5^80^. Therefore, we explored if ^2^H and ^15^N labeling could improve the spin-relaxation properties of semiquinone radicals and resolve additional distances by 4P-DEER. To the best of our knowledge, no studies on the enhancement of spin relaxation by isotopic labeling have been conducted with flavin semiquinone radicals, particularly in cellular environments. Indeed, the effect of the reducing environment of the *E. coli* cytoplasm on spin relaxation properties is poorly understood. We expressed Aer in *E. coli* grown in three fully isotopically substituted media: (^2^H/^14^N), (^1^H/^15^N), and (^2^H/^15^N) and characterized the relaxation properties of the flavin ASQ.

The cw-ESR spectra of Aer expressed in the four (^1^H/^14^N, ^1^H/^15^N, ^2^H/^14^N, ^2^H/^15^N) isotopically defined media was immediately revealing (**Figure 4a**). Under all labeling conditions, g-factors of 2.004 ± 0.001 (measured by comparison to a 2,2-diphenyl-1-picrylhydrazyl (DPPH) standard (g_DPPH_ = 2.0036)), were insensitive and in good agreement with previously reported g-factors for the semiquinone radical state in flavoproteins^13,81^. Relatively complex hyperfine interactions are expected for the flavin radical because the isoalloxazine ring contains multiple hydrogen and nitrogen atoms. However, these hyperfine peaks are typically not well resolved in ^1^H/^14^N and ^1^H/^15^N conditions due to the ESR linewidth. ENDOR experiments can help identify the ^1^H hyperfine coupling interactions (**Figure 2b**); nonetheless, the contributions of the nitrogen nuclei are usually obscured even in ENDOR. In the samples of Aer grown in deuterated media (^2^H/^14^N, ^2^H/^15^N), the proton hyperfine interactions on the flavin ring were suppressed and the nitrogen hyperfine interactions became apparent. The cw-ESR spectra indicated that the underlying nitrogen hyperfine peaks can be attributed to two major nitrogen atoms - N5 and N10^14,82^. For the (^2^H/^14^N) Aer sample, we observed a characteristic nine-line cw-ESR spectra (2 x I + 1)(2 x I +1) = 9, corresponding to the interaction of two non-equivalent ^14^N atoms (I = 1). For the (^2^H/^15^N) Aer sample, we observed four-line spectra (2 x I + 1)(2 x I +1) = 4 corresponding to the interaction of two non-equivalent ^15^N nuclei (I = 1/2). The spectral lineshapes of the cw-ESR spectra were simulated using the EasySpin toolbox in MATLAB^83^ with a consistent set of global parameters (**Figure S4**). In addition, hfccs (**Table S3**) estimated for the two nitrogen atoms from 3P-ESEEM measurements carried out at Q-band (**Figure S5**) agree with the values obtained from cw-ESR measurements. These simulations were carried out in the rigid limit and the best fit values (**Table S3**) are close to the reported values for flavin radicals found by high-field ESR measurements^84^ and for FMN radicals immobilized in an agarose matrix^15^. Our measurements confirmed that the flavin moieties are near the rigid limit and thus fixed within the protein pocket.

**Figure 4:**
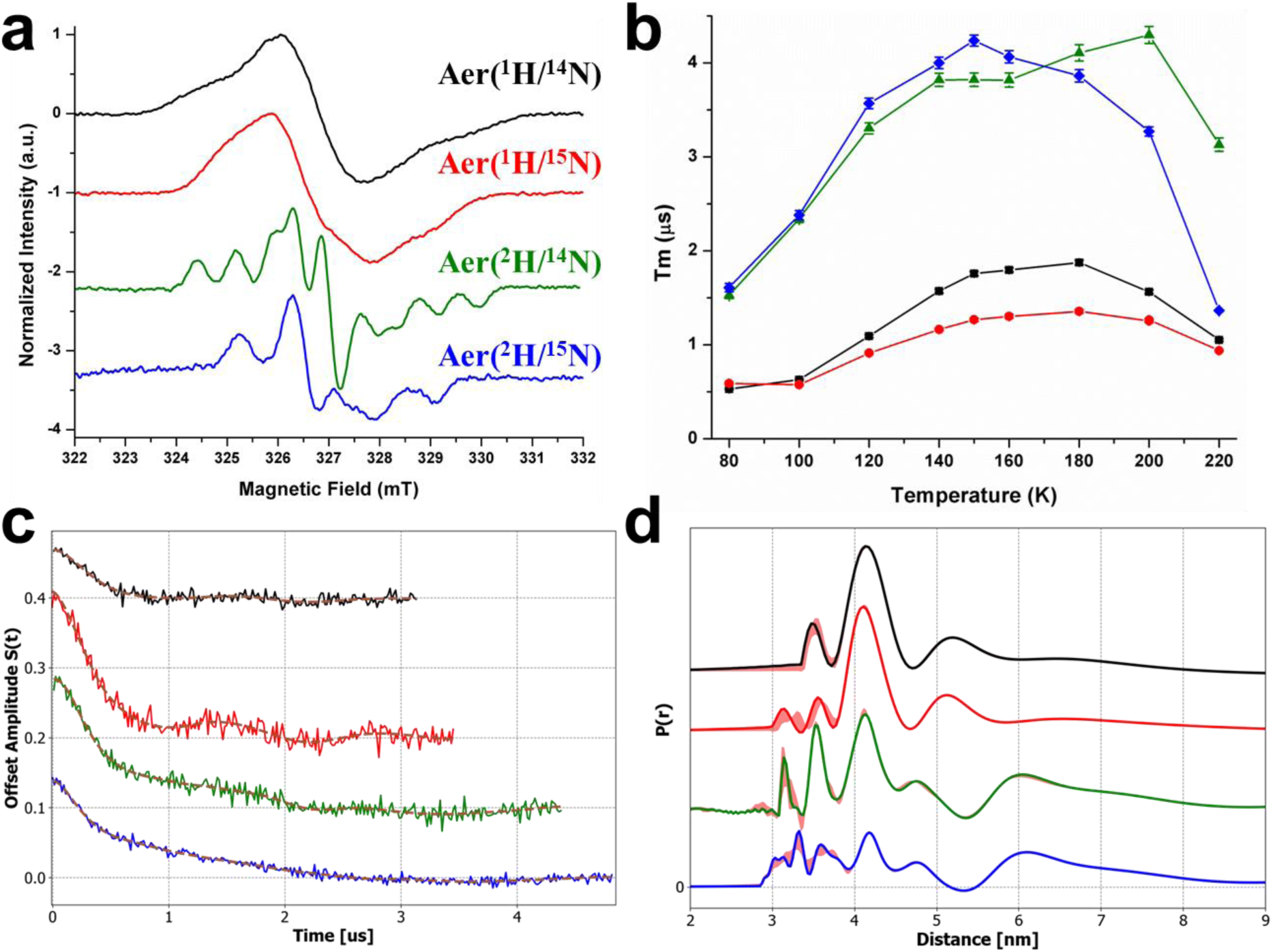
cw-ESR and pulsed ESR analysis of Aer co-expressed with CheA-CheW in cells grown in different isotopically enriched media: in normal LB media (black curves), in ^1^H/^15^N Celtone media (red curves) in ^2^H/^14^N Celtone media (green curves) in ^2^H/^15^N Celtone media (blue curves). (a) X-band *in cell* cw-ESR spectra of Aer measured at 293 K; (b) Phase memory relaxation time (T_m_) measured by Q-Band two pulses echo decay sequence (c) Time domain and reconstructed time domain (SF-SVD method^76^) (dashed brown line) measured at T = 150 K by Q-Band 4P-DEER sequence and (d) its associated distance distribution. Errors in the distance distributions represented by red shading and calculated as described in Srivastava et al.^76^.

Next, we investigated the effect of isotopic modifications on the phase memory time T_m_ (**Figure 4b**) and on the spin lattice relaxation time T_1_ (**Figure S6**) of the flavin radical *in cell*. Earlier studies have characterized the temperature dependence on T_1_ and T_m_ for related semiquinone radicals^85,86^. In particular, the presence of methyl groups on isoalloxazine ring promotes additional spin relaxation at lower temperatures owing to the modulation of hyperfine interactions by their rotation. T_m_ for all of the samples followed the same trend of a bell-shaped^86–89^ curve with a maximum between 150-180 K (**Figure 4b**). Deuteration of the entire protein enhanced the T_m_ by a factor of ∼3 at the T_m_ maxima whereas ^15^N labeling produced only minor improvements. These bell-shaped curves derive from two phenomena: below 120 K, the magnetic inequivalence of hindered methyl protons (H7, H8) leads to additional cross-relaxation, whereas above 180 K, molecular motions increases and produces faster spin dephasing rates^90^.

The spin lattice relaxation time T_1_ decreased monotonically with the temperature, with values ranging from 0.2 ms (240 K) to values slightly greater than 1 ms at cryogenic temperatures (100 K and lower) for unlabeled Aer (^1^H/^14^N) (**Figure S6**). Isotopic replacement of ^14^N with ^15^N has marginal effects on T_1_, whereas deuterium labeling enhanced the T_1_ by a factor ∼2-4, depending on the temperature. ^2^H MIMS ENDOR measurements confirmed deuterium substitution did not alter the flavin electronic structure (**Figure S7 and Table S4**). Interestingly, unlike nitroxide spin labels^87^, exchange of the protonated protein into a deuterated buffer alone does not enhance the spin relaxation of flavins significantly and no ^2^H-ENDOR signal was detected in the sample exchanged with deuterated buffer (**Figure S8**). This lack of substitution likely confirms that the FAD cofactor, rigidly bound in the PAS domain of Aer, is inaccessible to the solvent.

Next, we performed 4P-DEER on the isotopically labeled *in cell* Aer/CheA/CheW sample, with the goal of probing longer distances with longer evolution times. Thus, isotopically labeled AerCheACheW ^1^H/^14^N, ^1^H/^15^N, ^2^H/^14^N and the ^2^H/^15^N samples were collected until 3.1, 3.5, 4.4 and 4.9 µs evolution times, respectively (**Figure 4c** and **Table S5**).

A clear dipolar oscillation is observed for all of the samples with a period of ∼ 1.3 µs corresponding to the homodimer flavin separation of ∼ 4.1 nm. (**Figure 4d**). These results indicate that isotopic labeling did not significantly alter the arrangement of the PAS domains in Aer. Interestingly, additional distances from 3.2 nm to 6.5 nm were observed in all samples. We stress again that longer distances greater than 6 nm should be viewed with caution because evolution times were limited to a maximum of 5 μs. Nonetheless, these additional distances *in cell* correspond with what would be expected for a TODs assembly state within a higher array organization^29–33,61^ Modeling of the Aer oligomeric assembly based on how other chemoreceptors associate into hexagonal arrays^29–33,61^ (**Figure 5a**) reveals that the separation of the flavin centers in the assembly will only depend on a few parameters. The vertical position of the flavin relative to the membrane should be the same for each receptor due to the positioning of the transmembrane helices and known structures of the PAS domain^28^. Hence the flavin separations will largely be determined in the 2 dimensions parallel to the membrane. Furthermore, the conserved interactions of CheW and CheA with the membrane-distal receptor tips should provide a similar TODs arrangement as observed for other receptor arrays^29–33,61,91^. Two trimers associate in “core-complexes” around one dimeric CheA^91,92^, and then these core complexes can be extended into hexagonal arrangements by rings formed from CheA P5 and CheW. The higher-order contacts between core complexes could also produce close distances among the flavin centers, but these interactions are somewhat uncertain as small changes at the CheA/CheW baseplate could propagate to much larger shifts at the level of the PAS domains, some 19 nm away. Nonetheless, the inter-dimer flavin distances will primarily depend upon the rotation angle of the flavin-to-flavin vector relative to a vector between the dimer center and the trimer center (α,β,γ, **Figure 5b,c**). For example, with a fixed α=β=γ angle plane geometry (**Table S6**), the distances of the *in cell* distributions of the isotopically labeled AerCheACheW (^2^H/^14^N) and (^2^H/^15^N) at 3.6 nm and 4.7 nm could arise from a TODs with fixed angles of ∼120° (**Table S6 and Figure 5a**). A TODs with an angle of α=β=γ=120° would induce two additional longer distances at 7.2 nm and 7.7 nm that could explain the large peaks observed in the 7 nm region. Nevertheless, this simple geometry model of a TODs fails to explain the 3.2-3.3 nm distance peaks.

**Figure 5:**
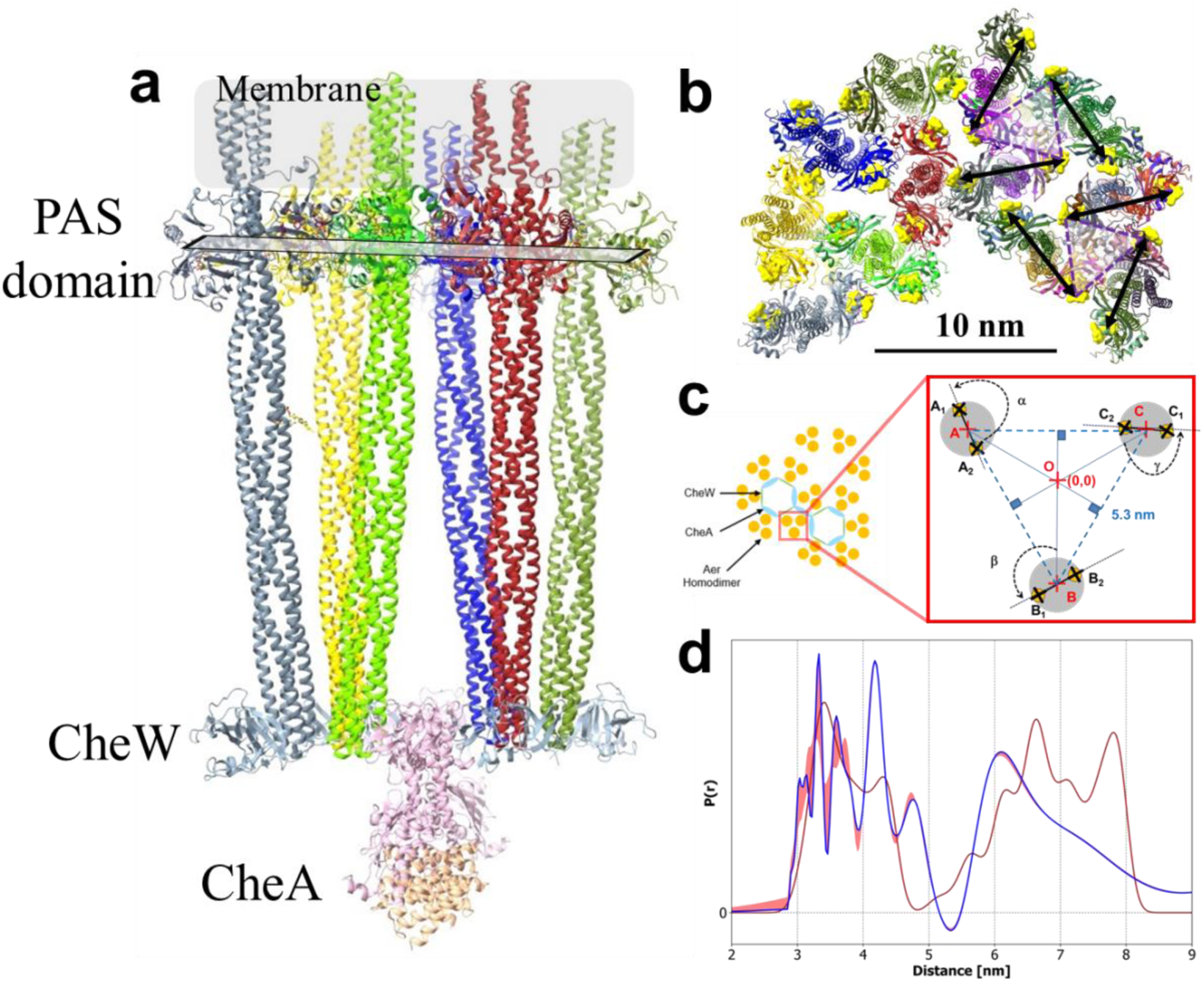
Aer forms higher-order structures *in cell*. (a) Model of a trimer of dimers (TODs) of AerCheACheW carried out with ChimeraX software^93^. The reconstruction is based on an alphafold^94^ reconstruction of Aer/CheA/CheW homodimer. The organization is showing that angles (α,β,γ) between 115° to 125° are required to reproduce a TODs without the PAS domains clashing. (b) Normal view of the reconstruction of four juxtaposed TODs of AerCheACheW forming “two core complexes”. Yellow surfaces represent the solvent-excluded surfaces of the FAD center in the PAS domain. (c) Schematic representing the larger organization of TODs within the hexagonal arrays. In the inset, yellow dots with black crosses represent the FAD center in the PAS domain. The two-fixed distances in this model are the 4.1 nm separating the PAS domain in a homodimer, and the 5.3 nm distance prescribed by a Tar/Tsr chemosensor array observed by cryo-electron tomography in minicells^29^. The angle α, β, γ used for calculating distances within a TODs induced a set of four distances between the FAD (**Table S6**). (d) Comparison of a predicted distance distribution (brown curve) obtained from the homology model (**Figure 5b and Table S7**) and an experimental distance distribution obtained for Aer co-expressed with CheA-CheW in Celtone ^2^H/^15^N media (blue curve).

We hypothesize that those peaks could derive from flavin-to-flavin distances that arise from two or more TODs (**Figure 5b**). To explore this possibility, distances between every flavin (N_5_ to N_5_ distances) were cataloged in an array model that contained 2 core complexes (4 TODs) related by hexagonal symmetry. Distances larger than 8 nm were excluded because they would not be detected by pulse dipolar spectroscopy (**Table S7**). A distance distribution was generated by summing single gaussian curve for every distance with a standard deviation of 0.15 nm. The predicted distance distribution compares well to the experimental distance distribution obtained with the isotopically labeled AerCheACheW ^2^H/^15^N (**Figure 5d)** with multiple distances around 3.2-3.6 nm and around 6-7 nm. No distances are observed below 3 nm due to the excluding radius of the PAS domain. We note that the 5.2 nm peaks observed in some spectrum, as well as the fine structure in the experimental distributions, could derive from other factors that include: a non-identical angle geometry (α≠β≠χ), a membrane curvature effect that alters planarity or conformational sampling, the latter due to heterogeneity of assembly or even the redox state of the Aer homodimer. Furthermore, the weighting of the distances in the model histogram will depend on the relative amounts of TODs, core complexes and larger assemblies *in cell*, factors that are difficult to predict.

Distances indicative of higher-order structures with Aer were not observed *in vitro* because detergent solubilization breaks down TODs into homodimers and it is very challenging to incorporate homogeneous, aligned TODs into nanodiscs in sufficient yields to carry out PDS. Furthermore, the extended structures of the arrays are more difficult to produce *in vitro* and have only been assembled with fragments of chemoreceptors^91,95–99^. Our results highlight the power of using isotopic labeling and *in cell* ESR to probe these large macromolecular complexes in their native environment.

### Flavoprotein Probes for *in cell* ESR

To expand *in cell* ESR spectroscopy measurements to proteins lacking a native flavin cofactor, we used a small light-oxygen-voltage (LOV) sensing protein containing FMN cofactor as a spin probe^100,101^. Our previous work had shown that LOV domains lacking the adduct forming cysteine residue produce stable radicals upon photoreduction^100,102^.

One of the major challenges for structural *in cell* studies involves the intracellular delivery of the spin probe^103^. First, the spin label or spin-labeled protein has to cross the outer membrane, cell wall and inner membrane, which is usually achieved with semi-destructive techniques such electroporation, osmotic pressure or microinjection. If the spin label is delivered separately it then must be specifically targeted to the protein of interest, a challenging endeavor. To overcome these issues, we genetically fused iLOV^101^ (12.4 kDa) to the N-terminal P1 domain of the full-length histidine kinase CheA (77 kDa). CheA is composed of five domains, P1-P5 (**Figure 1c,d**). The P1-domain contains the histidine residue that normally undergoes phosphorylation, the P2 domain constitutes the secondary messenger (CheY) binding site, the P3 domain dimerizes CheA, the P4 domain contains the ATP binding site and the P5 domain interacts with the chemoreceptors (Aer/Tar) and CheW. The P1 and P2 domains of CheA are highly dynamic and are connected by the flexible linkers (L1 and L2)^97^. Recent evidence suggests that the P1-domains of CheA may dimerize and associate with the core P3-P4-P5 domains in the inhibited off-state of the kinase when it is bound to chemoreceptors^30,91^.

First, we confirmed that the attachment of iLOV does not affect the autophosphorylation activity of CheA *in vitro* (**Figure S9**). Second, we generated a NSQ radical in iLOV by using blue light irradiation as previously characterized^104,105^. The concentration (**Table S8**) was evaluated by cw-ESR and the nature of the radical *in cell* was confirmed by pulsed ^1^H-ENDOR spectroscopy (**Figure S10**).Third, we checked the stability of the radical in cellular condition (**Figure S11**) by cw-ESR. The photogenerated radical was found to persist for hours with a half time t_1/2_ ∼ 5 h with a mono-exponential decay time constant t_decay_ = 7.4 ± 0.4 h.

We then investigated the influence of the cell environment on the conformation of the P1 domain in CheA. The 4P-DEER spectrum (**Figure 6**) of iLOV-CheA in solution revealed little or no dipolar signal; the weak, long distance observed around 4-4.5 nm is likely an artefact due to the background subtraction. We then measured CheA-iLOV *in cell* at concentrations similar to those measured *in vitro* (**Table S8**). In contrast to the *in vitro* sample, 4P-DEER experiments on CheA-iLOV either alone or co-expressed with CheW and Tar receptor indicated that the iLOV domains on P1 domain of CheA reside close to each other, producing a dipolar interaction of 2.8 nm (which corresponds to an oscillation period of 420 ns) (**Figure 6)**.

**Figure 6:**
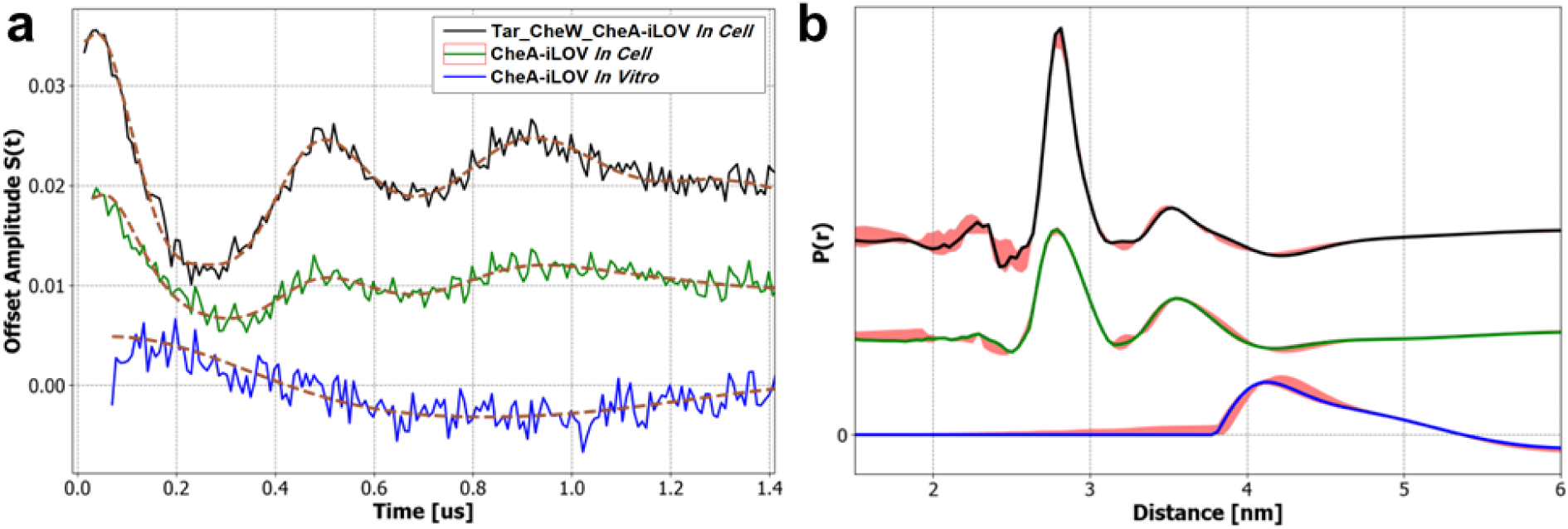
Q-Band 4PDEER measurements of CheA-iLOV in BL21 *E. coli* co-expressed with Tar and CheW (black curve) or alone (green curve) and of purified CheA-iLOV (blue curve). (a) Time domain and reconstructed time domain (SF-SVD method^76^) (dash brow line) measured at T = 180 K and (b) its associated domain distribution P(r). Errors in the distance distributions represented by red shading and calculated as described in Srivastava et al.^76^.

The modulation depths for these experiments were relatively small (λ = 4.3 % for Tar-CheW-CheA-iLOV *in cell* versus 1.3 % and 0.4 % for CheA-iLOV *in cell* and *in vitro*, respectively) (**Table S8**) The comparison of CheA-iLOV *in vitro* to *in cell* indicates that the cellular environment alone favors some interaction of the CheA P1 domains, perhaps due to molecular crowding in the cytoplasm. However, despite similar expression levels, when CheA-iLOV was co-expressed with its partners, the signal from the iLOV-iLOV interaction increased substantially.

Studies on Tar-CheA-CheW complexes indicate that association of CheA with CheW and receptors facilitates P1-P1 interactions, which would thereby bring the iLOV domains into close proximity of each other^91,97,106,107^. When bound to receptors CheA may adopt different conformations associated with so-called kinase-on and kinase-off activity states^91,106,107^. Either or both of these states may involve P1 dimerization^91,106,107^. Efforts to alter the kinase activity state of CheA by adding the Tar attractant ligand aspartate or repellant Ni^2+^ produced little difference in the distance domain (**Figure S12**), which could indicate that either the P1 domains remain dimerized regardless of activity state, or that ligand-induced shifts in the activity state involve a population of molecules that is too small to observe under these conditions. Nonetheless, the CheA-iLOV fusion successfully reported on the conformational state of CheA within receptor arrays and revealed domain arrangements that are difficult to recapitulate in reconstituted systems and are generally not assumed by the isolated kinase.

## CONCLUSIONS

New methods to interrogate the structural and biophysical properties of proteins in their native cellular environments are in demand. Flavin cofactors with their ability to stabilize radical states have the potential to serve as genetically encodable probes for ESR spectroscopy. Here we realize this concept by investigating the behavior of the FAD-binding aerotaxis receptor Aer in its native signaling assembly and by developing a small-flavoprotein probe iLOV that can be fused to any protein of interest. As Aer receptor conformations and oligomeric states are difficult to reproduce *in vitro*, *in cell* ESR spectroscopy provided new insight into its assembly state with CheA and CheW. Q-band ^1^H Davies ENDOR measurements revealed that Aer primarily forms an anionic semiquinone radical *in cell* with hfccs consistent with other ASQ-forming flavoproteins. This redox state has important implications for the Aer signaling mechanism and comments on the low potential of its membrane environment^28^. Q-Band 4P-DEER measurements revealed a flavin-flavin distance of 4.1 nm between Aer subunits within the homodimer, which agrees well with previous *in vitro* measurements of isolated, chemically-reduced Aer^27^. In comparison to purified receptors, the *in cell* PDS measurements revealed additional distances consistent with chemoreceptor array formation in the cytoplasmic membrane. Additionally, temperature screening and isotopic labeling with ^2^H and ^15^N was performed to optimize relaxation properties for *in cell* PDS measurements, which enabled visualization of signaling complexes difficult to observe *in vitro*. Moreover, we extend the approach to a general target protein by incorporating a small flavin-containing domain, wherein stable radicals can be induced with light directly *in cell*. In particular, CheA-iLOV when expressed in cells with its signaling partners gave dipolar signals characteristic of domain associations that are not observed with purified samples, thereby demonstrating the importance of *in cell* measurement and the means to expand this method to a wide range of applications. Overall, this study should serve as a benchmark for future *in cell* investigations of flavoproteins.

## MATERIALS AND METHODS

### Constructs and protein expression in *E. coli* BL21 (DE3)

Full length *E. coli* Aer (1-506), Aer mutants (AerC193HC203H and AerY93HC193HC203H) were cloned into the pET-28a plasmid. Tar (1-553) was either cloned in pET-28a for the Tar-CheA-CheW construct or cloned into pCDF-DuetI for the Tar-CheW-CheA-iLOV construct. The histidine kinase CheA was cloned into the first multiple cloning sites (MCS) site of a pACYC-DuetI vector using the NcoI/BamHI restrictions sites whereas the coupling protein CheW was cloned into the second MCS site of pACYC-DuetI using the NdeI/XhoI sites. Aer along with its mutants in pET-28a and CheA/CheW in pACYC-DuetI were co-transformed into BL21 and selected on agar plates against kanamycin and chloramphenicol.

CheA-iLOV was constructed by cutting the CheA/CheW coding sequence from the pACYC-DuetI vector with NcoI/XhoI and inserting it into pET-28a-iLOVf (Addgene plasmid # 63723) cut at the same restrictions sites to produce the fused CheA-iLOV in MCS-1 and CheW in MCS-II. CheA-iLOV/CheW in pET-28a was co-transformed with Tar in pCDF-DuetI and selected on agar plates containing kanamycin and streptomycin. CheA-iLOV expressed in BL21-DE3 cells were purified by nickel affinity and size exclusion chromatography on a preparative Superdex^TM^ 200 pg HiLoad 26/600 column (**Figure S9**).

#### Protein purification

Plated cells with corresponding plasmids fragment and antibiotics (Aer (pET-28a) with coexpression of CheA/CheW (pACYC) and CheA-iLOV (pET-28a)) were grown in a 2x2L LB in 2x6L vials at 37°C for around 6h and induced at an OD=0.6 with 1mM IPTG. The vials were then shaken overnight at 17°C. The cell pellets were harvested by centrifugation at 10000 rpm for 60 min at 4°C. Cell pellets were resuspended in Lysis buffer (50 mM Tris, 200 mM KCl, 10 % glycerol, pH = 8.0) and was digested with 60 mg Lysozyme and 0.1 mM PMSF for an hour at 4°C. The Emulsiflex-C3 was equilibrated with lysis buffer and used to lyse the cell suspension at 15,000-18,000 psi in 3 passes.

##### Purification of Aer

As Aer is a membrane protein an initial low-spin centrifugation of the cell lysate was carried out at 20,000 rpm for 45 min at 4 °C to remove cell debris. The low-spin supernatant was then ultracentrifuged at 100,000 x g for 1 h at 4°C to acquire the insoluble fraction. The insoluble fraction was resuspended in lysis buffer at a 1:2 ratio and the insoluble Aer protein was solubilized overnight at 4 °C with 1 % Lauryl Maltose Neopentyl Glycol (Anatrace). The solubilized membrane fraction was centrifuged at 4,000 RPM for 10 min at 4°C to remove any additional insoluble fraction. The supernatant was mixed with 0.1 mg/ml of FAD then was gently rocked with 5 mL bed volume of lysis buffer prewashed Nickel NTA resin (Nickel NTA HTC Agarose Resin from GoldBio) overnight at 4°C. The next morning, the resin was washed three times with lysis buffer containing 0.1 % LMNG. The protein was then eluted from the resin with the elution buffer (50 mM Tris, pH = 8.0, 150mM NaCl; 10 % glycerol, 200 mM imidazole, pH = 8.0) and concentrated to the highest concentration possible with a 15 mL 50 kDa cutoff amicon filter and washed two times with the resuspension buffer (25 mM Tris, pH = 8.0, 150mM NaCl; 10 % glycerol, 1 % LMNG, pH = 8.0). Aer protein was directly used for analysis after purification.

##### Purification of CheA-iLOV

The cell lysate was centrifugated at 20,000 rpm for 45 min at 4°C. The yellow supernatant was directly mixed with 5 mL bed volume of lysis buffer prewashed Nickel NTA resin (Nickel NTA HTC Agarose Resin from GoldBio) and left gently rocking overnight at 4°C. The next morning, the resin was washed three times with lysis buffer. The protein was then eluted from the resin with the elution buffer (50 mM Tris, pH = 8.0, 150mM NaCl; 10 % glycerol, 200 mM imidazole) and concentrated to the highest concentration possible with a 15 mL 50 kDa cutoff amicon filter. CheA-iLOV was stored at 4°C for future analysis.

#### CheA autophosphorylation assays

CheA autophosphorylation was monitored by ^32^P incorporation. All radioassays were carried out in 5 mM Tris (pH = 7.5), 50 mM KCl, 10 mM MgCl_2_, 0.5 mM DTT, 0.5 mM EDTA. Samples were prepared with 2 µM CheA and 4 µM CheW in a final volume of 25 µL. Following incubation for ∼15 min, [γ-^32^P]ATP was added to a final concentration of 1 mM and the reaction was quenched after 30 seconds using 4x SDS-PAGE loading buffer containing 50 mM EDTA at pH = 8.0. The samples were loaded onto a 4-20 % Tris-glycine polyacrylamide protein gel purchased from Invitrogen. Gel electrophoresis was carried out for 35-60 minutes at 125 V constant voltage using a Tris-gly-SDS running buffer. The resulting gels were dried in a Bio-Rad Gel Dryer overnight and placed in a radiocassette for >20 hours prior to imaging on a Typhoon Image Scanner.

### Preparation of *in cell* ESR samples

All the proteins were co-expressed with the appropriate antibiotic in *E. coli* BL21(DE3) cells under 1 mM Isopropyl β-d-1-thiogalactopyranoside (IPTG) induction. For the natural isotope experiments (^1^H and ^14^N), the cells were grown in autoclaved freshly prepared Luria broth media while the appropriate Celtone complete media (^2^H, 97 % or ^15^N, 98 % or ^2^H, 97 % and ^15^N, 98 %) from Cambridge Isotope Laboratory® was used for the isotope enriched experiments. Cells were grown in the appropriate media to an O.D. of ∼ 0.5-0.6 and induced for 3 h at 37 °C (protonated media) or overnight (∼ 16 h) at room temperature (deuterated media). The cells where then spun down, washed and resuspended in a small amount of 25 % glycerol (25 % d_3_-glycerol in D_2_O for the deuterated samples). The cells were then transferred into a ∼ 1 mm I.D. capillary tube (Kimble® 71900-50 KIMAX® 50 µL precision microcapillaries) and spun down with a hematocrit centrifuge. The iLOV protein was irradiated at room temperature with a λ = 457 ± 10 nm Blue LED (LZ1-00B202 from LedEngin) for 3 minutes through the window of the ER 4123SHQE Bruker cavity to generate the semiquinone radical (see **Figure S10** for the cw-ESR spectra). The X-band cw-ESR measurements were recorded at room temperature and the samples were plunge frozen in liquid N_2_ before carrying out the Q-band pulsed ESR measurements.

### Preparation of *in vitro* ESR samples

30 μM Aer purified protein stock solution (dissolved in 1 % LMNG detergent) was supplemented with 25 % glycerol and 10 mM DTT. 20 μL of this solution was quickly inserted into the ESR capillaries and any oxygen was purged by 3 freeze-pump-thaw cycles with argon. The flame-sealed capillary was irradiated with λ = 457 ± 10 nm Blue LED (LZ1-00B202 from LedEngin) for 20 min. The irradiated sample was followed *in situ* by cw-ESR and the sample was frozen just right after the maximum of intensity was generated (30 μM concentration in spin).

250 μM of CheA-iLOV purified proteins stock solution was supplemented with 25 % glycerol and 10 mM DTT. 20 μL of this solution was quickly inserted in the ESR capillaries and any oxygen was purged by 3 freeze-pump-thaw cycles with argon. The flame-sealed capillary was irradiated with λ = 457 ± 10 nm Blue LED (LZ1-00B202 from LedEngin) during 20 min. The irradiated sample was followed *in situ* by cw-ESR and the sample was frozen after the maximum of radical intensity was obtained (210 μM concentration in spin).

#### X-band cw-ESR

ESR spectra were recorded using a continuous wave X-band Bruker ElexSys E500 ESR spectrometer equipped with a ER 4123SHQE Bruker cavity. The acquisition parameters were fixed to 15 dB (6.325 mW) microwave power, 60 dB receiver gain, and 100 kHz modulation frequency. All cw-ESR spectra were recorded at 298 K with an amplitude modulation of 1 G to observe the hyperfine splitting. All of the simulations used to obtain the hyperfine coupling constants (hfccs) were carried out with EasySpin^83^ (6.0.6) as implemented in MATLAB (R2021a; refer to code in the supplementary.)

#### Q-Band pulsed ESR

The Q-band pulsed ESR analysis was carried out with an Elexsys E580 spectrometer equipped with a 10 W solid state amplifier. All of the pulsed experiments were achieved in an EN 5107D2 Cavity, Q-Band ENDOR Pulsed ESR. A Bruker E-580 AWG Arbitrary Waveform Generator was used for the microwave pulse generation of the 4P-DEER, ENDOR and 3P-ESEEM sequences. Microwave pulses were generated by the Super-QFT-Upgrade Microwave Bridge for the relaxation time measurements. The temperature was varied by using an ER 4118HV-CF10-L FlexLine Cryogen-Free VT System. The pulse length (varying with the coupling range of the resonator and the microwave power used) was determined with a rabi nutation measurement π-t_2_-π/2-t-π-t-echo. All of the pulsed ESR experiment with Aer and iLOV *in cell* were conducted at temperatures of 150 K - 180 K to maximize the phase memory time T_m_, as assessed from the temperature profiles presented in **Figure 4b**.

##### Time relaxation measurements

The resonator was critically coupled, and the magnetic field was chosen to be at the maximum peak signal intensity. The typical length of the microwave pulse was around 16 ns and 32 ns for a π/2 and π pulse, respectively, in these conditions.

The electron spin–lattice relaxation times T_1_ were measured over the temperature range by an inversion recovery pulse sequence, π–t_2_–π/2–t–π–t–echo by varying t_2_. For each trace, 512 data points were collected with an appropriate time increment to ensure complete magnetization recovery. The trace was fitted by a bi-exponential model. The longer time decay T_1L_ was reported in the **Figure S6**. Phase-memory times T_m_ were measured over the temperature range by a two-pulse echo decay sequence, π/2–t–π–t–echo, while varying the t. The curves were fit by an exponential decay: I(t)=I_0_*exp(-2t/T_m_).

##### Pulsed ENDOR measurements

Pulsed Mims sequence π/2-tau-π/2-t1-rf pulse-t2-π/2-t-echo was employed to detect ^2^H deuterium ENDOR signal. The π/2 pulses were chosen to be as short as possible and to be non-selective. To avoid the tau blind spot effect, the Mims ENDOR spectra were accumulated with a tau value varying from 120 ns to 150 ns. The different ^2^H Mims spectra were summed over these tau values. The Davies sequence π-t1-rf pulse-t2-π/2-t-π-t-echo was employed for measuring ^1^H proton signal as the hfccs of some protons can be very large in the semiquinone radicals (10-30 MHz for some positions). The length of the selective microwave pulse was around 60 ns and 120 ns for a π/2 and π pulse, respectively.

For both sequences, 1) the resonator was critically coupled, 2) t_1_ and t_2_ were fixed to 1 μs to avoid any overlap between the microwave pulses and the radiofrequency (rf) pulse and 3) a 20 µs radiofrequency (rf) pulse was applied with a 150W Bruker RF amplifier.

##### 4P-DEER measurements

4-pulse DEER sequence π/2-t1-π-t1-π(pump)-t2-π-t2-echo was carried out at 150 K. The resonator was undercoupled to increase the microwave bandwidth. As a result, typical π/2 and π pulses of 18 ns and 36 ns were used. The time domain data was background subtracted and distance distributions were obtained by the SF-SVD method^76^. Data processing of the SF-SVD-based method was used through the “SVDReconstruction” software (https://denoising.cornell.edu/). The distance domain was normalized (total probability fixed to 1).

The python scripts used for data treatments are available at https://github.com/TChauvire/.

## ASSOCIATED CONTENT

**Supporting Information.** Supplementary tables reporting 4P-Deer data analysis, comparing hfccs values measured by cw-ESR, pulsed ENDOR and 3P-ESEEM, reporting distances obtained by homology model; and supplemental figures including cw-ESR analysis of IPTG-induced Aer, cw-ESR spectra of Y93 Aer mutants, 4P-DEER of Aer with and without CheA/CheW co-expression, cw-ESR fits of Aer expressed in isotopic media, 3P-ESEEM spectra, T_1_ temperature profiles, ^2^H-Mims ENDOR spectra, ^2^H buffer exchange experiment, SDS-Page gels of CheA-iLOV purification, cw-ESR spectra of photogenerated CheA-iLOV, *in cell* kinetics of degradation of CheA-iLOV, 4P-DEER analysis of TarCheWCheA-iLOV in minimal media, and supplementary Matlab scripts for the cw-ESR and 3P-ESEEM simulation. This material is available free of charge via the Internet.

Raw and processed data can be accessed freely on the Github pages: https://github.com/TChauvire/.

## AUTHOR INFORMATION

### Author Contributions

All authors have given approval to the final version of the manuscript.

### Funding Sources

This work was supported by grants from the National Science Foundation: MCB 2129729 to BRC; National Institutes of Health: R35GM122535 to BRC and R35GM148272 to JHF; and by the National Institute Of General Medical Sciences (NIGMS) of the National Institutes of Health (NIH) under Award Number R24GM146107 and P41GM103521 to JHF and BRC

### Notes

The authors declare no competing financial interest.

## ACKNOWLEDGMENT

We thank Curt R. Dunnam and Walt P. Ford for the assistance and the repairs of the pulsed ESR spectrometer, Dr. Zachary Maschmann for the preparation and the expression of Aer site-directed variants, and Dr. Madhur Srivastava for the help with the use of the SF-SVD software.

## ABBREVIATIONS

ESR: electron spin resonance
cw-ESR: continuous wave ESR
ENDOR: electron nuclear double resonance
PDS: pulsed dipolar spectroscopy
4P-DEER: Four-pulse Double Electron-Electron Resonance
ESEEM: Electron Spin Echo Envelope Modulation
hfccs: hyperfine coupling constants
SF-SVD: Srivastava-Freed singular value decomposition method
rf: radiofrequency
mw: microwave
T_1_: spin-lattice relaxation time
T_m_: phase-memory relaxation time
TODs: trimers-of-dimers
FAD: flavin adenine dinucleotide
FMN: flavin mononucleotide
ASQ: anionic semiquinone
NSQ: neutral semiquinone
Aer: aerotaxis
Tar: aspartate chemoreceptor
LOV: Light Oxygen Voltage
PAS: Per-Arnt-Sim
NADH: reduced nicotinamide adenine dinucleotide
ATP: Adenosine triphosphate
*E. coli*: Escherichia coli

## Notes

### Competing Interest Statement

The authors have declared no competing interest.

