## Supplementary Information for "Flavoproteins as native and genetically encoded spin probes for *in cell* ESR spectroscopy"

Supplementary Tables: ........ S2

Supplementary Figures: ........ S8

Figure S1: IPTG-induced expression of Aer measured by *in cell* cw-ESR. ........ S8

Figure S2: Cw-ESR spectra for Aer and Aer site-directed variants. ........ S9

Figure S3: *In cell* 4P-DEER analysis of Aer co-expressed with and without CheA/CheW. ........ S10

Figure S4: X-band cw-ESR spectra of Aer grown in isotopically enriched media ........ S11

Figure S5: Aer($^{1}$H/$^{14}$N) and Aer($^{1}$H/$^{15}$N) 3P-ESEEM fitting ........ S12

Figure S6: Temperature profiles of the spin lattice relaxation time $T_1$ of isotopically enriched Aer co-expressed with CheA and CheW *in cell*. ........ S13

Figure S7: Aer($^{2}$H/$^{14}$N) and Aer($^{2}$H/$^{15}$N) Mims $^{2}$H-ENDOR spectra ........ S14

Figure S8: *In cell* deuterium solvent exchange experiment with Aer($^{1}$H/$^{14}$N)-CheA-CheW........ S15

Figure S9. Expression and Activity of CheA-iLOV. ........ S16

Figure S10. Cw-ESR and $^{1}$H-ENDOR spectra of CheW-CheA-iLOV. ........ S17

Figure S11: Kinetic of degradation of CheA-iLOV in cellular environment. ........ S18

Figure S12: *In cell* 4P-DEER measurements of TarCheWCheA-iLOV in different growth conditions in minimal media ........ S19

MATLAB code for global fitting ........ S20

References: ........ S23

### Supplementary Tables:

**Table S1: 4P-DEER experimental parameters[1] of Aer co-expressed with or without CheA/CheW (*in cell* measurement) and Aer dissolved in LMNG detergent *in vitro*. (see Figure 3)** These parameters were obtained after zero-order phasing of the experimental data and background subtraction with the function: $B(t) = exp(a * t + b)$. The data were subtracted by this background correction as suggested by Ibáñez et. al.[2]. The time domain was truncated (reported as $t_{max}$) and was fixed to 19/20 of the global range of the time domain. The "Offset Time" (noted as $t_{min}$) was determined using the "correctzerotime.py" function of the Deerlab software[3]. The noise level was evaluated using the standard deviation of the imaginary part of the data divided by the maximum value of the real part. Signal to noise ratio was then deduced by the ratio of the modulation depth λ over the noise level σ. The modulation depth λ was evaluated by subtracting $S(t=t_{min}) - B(t=t_{min})$. Finally the background decay η was obtained by the relation $\eta = 1 - \frac{B(tmax)}{B(tmin)}$.

| | ***In cell* 4P-DEER** | | ***In vitro* 4P-DEER** |
|---|---|---|---|
| **Parameters :** | **Aer($^1$H/$^{14}$N) co-expressed with CheA-CheW** | **Aer($^1$H/$^{14}$N) without co-expression of CheA-CheW** | **Aer($^1$H/$^{14}$N) in detergent (LMNG)** |
| **$t_{max}$ (ns)** | 2330 | 2370 | 2330 |
| **Offset Time $t_{min}$ (ns)** | 2 | 2 | 2 |
| **Noise level σ** | 0.0061 | 0.0029 | 0.0093 |
| **Modulation depth λ** | 0.157 | 0.036 | 0.078 |
| **DEER SNR** | 25.9 | 12.4 | 8.4 |
| **Background Decay η** | 0.2818 | 0.0610 | 0.0178 |
| **Background parameters (a[ns$^{-1}$],b)** | -0.1840<br>$-1.4 \cdot 10^{-4}$ | -0.043<br>$-2.7 \cdot 10^{-5}$ | -0.0888<br>$-2.7 \cdot 10^{-5}$ |
| **RMSE (Background subtraction)** | 0.0570 | 0.0155 | 0.0311 |
| **Spin concentration measured by cw-ESR (µM)** | 32 | 35 | 29 |

**Table S2: 4P-DEER experimental parameters[1] of Aer co-expressed with or without CheA/CheW. (see Figure S3)** These parameters were obtained after zero-order phasing of the experimental data and background subtraction with the function: $B(t) = exp(a * t + b)$. The data was then subtracted by this background correction as suggested by Ibáñez et. al.[2]. The time domain was truncated (reported as $t_{max}$) and was fixed to 79/80 of the global range of the time domain. The "Offset Time" (noted as $t_{min}$) was determined using the "correctzerotime.py" function of the Deerlab software[3]. The noise level was evaluated using the standard deviation of the imaginary part of the data divided by the maximum value of the real part. Signal to noise ratio was then deduced by the ratio of the modulation depth $\lambda$ over the noise level $\sigma$. The modulation depth $\lambda$ was evaluated by subtracting $S(t=t_{min}) - B(t=t_{min})$. Finally the background decay $\eta$ is obtained by the relation $\eta = 1 - \frac{B(tmax)}{B(tmin)}$.

| | Aer($^1H/^{14}N$) co-expressed with CheA and CheW | | | | Aer($^1H/^{14}N$) without co-expression of CheA and CheW | | |
|---|---|---|---|---|---|---|---|
| **Parameters :** | **1st** | **2nd Repeat** | **3rd Repeat** | **4th Repeat** | **1st** | **2nd Repeat** | **3rd Repeat** |
| **$t_{max}$ (ns)** | 2426 | 3146 | 2894 | 3406 | 2370 | 2442 | 2954 |
| **Offset Time $t_{min}$ (ns)** | 2 | -6 | -2 | -2 | 2 | -6 | -6 |
| **Noise level $\sigma$** | 0.006 | 0.006 | 0.011 | 0.022 | 0.003 | 0.009 | 0.012 |
| **Modulation depth $\lambda$** | 0.157 | 0.049 | 0.092 | 0.092 | 0.036 | 0.032 | 0.031 |
| **DEER SNR** | 25.9 | 7.9 | 8.5 | 4.2 | 12.4 | 3.6 | 2.7 |
| **Background Decay $\eta$** | 0.29 | 0.09 | 0.26 | 0.31 | 0.06 | -0.04 | -0.05 |
| **Background parameters (a[$ns^{-1}$],b)** | -0.184 $-1.4\cdot10^{-4}$ | -0.070 $-3.1\cdot10^{-5}$ | -0.122 $-1.1\cdot10^{-4}$ | -0.111 $-1.1\cdot10^{-4}$ | -0.043 -2.7·10-5 | -0.094 $+1.6\cdot10^{5}$ | -0.083, $+1.5\cdot10^{5}$ |
| **RMSE (Background subtraction)** | 0.057 | 0.045 | 0.032 | 0.030 | 0.015 | 0.072 | 0.091 |
| **Spin concentration measured by cw-ESR (µM)** | 32 | 32 | 26 | 26 | 35 | 11 | 11 |

**Table S3: Hyperfine coupling constants (hfccs) of N5, N10 of the flavin cofactor obtained after fit of the X-band cw-ESR spectra (see Figure S4) and the Q-band 3P-ESEEM spectra (Figure S5).** The MATLAB scripts used for fitting are reported in the supplementary information.

| | **cw-ESR (Global)** $A_{xx}$ $A_{yy}$ $A_{zz}$ **(MHz)** | **3P-ESEEM: $^1H/^{14}N$** $A_{xx}$ $A_{yy}$ $A_{zz}$ **(MHz)** | **3P-ESEEM: $^1H/^{15}N$** $A_{xx}$ $A_{yy}$ $A_{zz}$ **(MHz)** |
|---|---|---|---|
| **N5** | 1.7, 1.2, 56.3, (2.4), (1.7), (79.0) | 1.3, -0.8, 58.6 | 2.4, 0.08, 60.6, (3.4), (0.1), (84.9)[(a)] |
| **N10** | 4.5, 4.4, 23.1, (6.3), (6.2), (32.4) | 3.6, 4.1, 21.3 | 2.3, 2.1, 26.2, (3.3), (2.9), (36.7)[(a)] |

(a) Values in parentheses are conversion of $^{15}N$ hfcc to an equivalent $^{14}N$ hfcc via the gyromagnetic ratio $|\gamma_{15N}/\gamma_{14N}|$=1.4028.

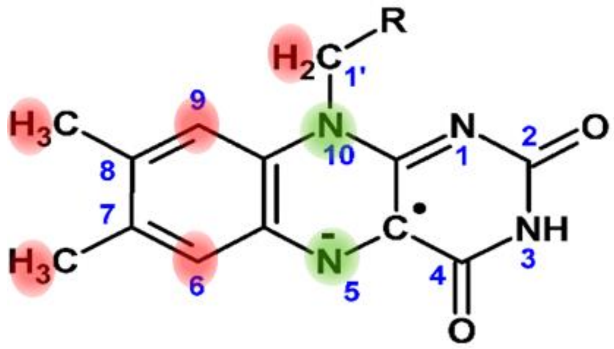

**Table S4: Hyperfine Coupling Constants (hfccs) in MHz of the six protons of the flavin cofactor after analysis by Q-band pulsed ENDOR (Figure 3 and Figure S7).**

| **Sample:** | **$^1H/^{14}N$** | **$^1H/^{15}N$** | **$^2H/^{14}N$** | **$^2H/^{15}N$** |
|---|---|---|---|---|
| **Pulsed Experiments:** | **$^1$H Davies Endor** | | **$^2$H Mims Endor** | |
| **$^1$H(C9)** (MHz) | 0.88 | 1.03 | - | - |
| **$^1$H(C1'b)** (MHz) | 1.81 | 1.76 | - | - |
| **$^1$H(C7a)** (MHz) | 3.47 | 3.47 | 0.59 (3.82)[(a)] | 0.59 (3.82)[(a)] |
| **$^1$H(C6)** (MHz) | 7.97 | 7.72 | 1.21 (7.90)[(a)] | 1.19 (7.77)[(a)] |
| **$^1$H(C1'a)** (MHz) | 8.70 | 8.65 | - | - |
| **$^1$H(C8a)** (MHz) | 10.36 | 10.36 | 1.60 (10.44)[(a)] | 1.60 (10.44)[(a)] |

[(a)] Values in parentheses are a conversion of $^2H$ hfcc to $^1H$ hfcc via the gyromagnetic ratio $|\gamma_{1H}/\gamma_{2H}|$=6.516.

**Table S5: 4P-DEER experimental parameters[1] of Aer co-expressed with CheA/CheW in rich media of different isotopic compositions (see Figure 4).** These parameters were obtained after zero-order phasing of the experimental data and background subtraction with the function: $B(t) = exp(a * t + b)$. The data were then subtracted by this background correction as suggested by Ibáñez et. al.[2]. The time domain was truncated (reported as $t_{max}$). The "Offset Time" (noted as $t_{min}$) was determined using the "correctzerotime.py" function of the Deerlab software[3]. The noise level was evaluated using the standard deviation of the imaginary part of the data divided by the maximum value of the real part. Signal to noise ratio was then deduced by the ratio of the modulation depth λ over the noise level σ. The modulation depth λ was evaluated by subtracting S(t=$t_{min}$) - B(t=$t_{min}$). Finally the background decay η is obtained by the relation $\eta = 1 - \frac{B(tmax)}{B(tmin)}$.

| Parameters : | Aer($^1H/^{14}N$) | Aer($^1H/^{15}N$) | Aer($^2H/^{14}N$) | Aer($^2H/^{15}N$) |
|---|---|---|---|---|
| **$t_{max}$ (ns)** | 3146 (79/80) | 3466 (79/80) | 4394 (19/20) | 4858 (79/80) |
| **Offset Time $t_{min}$ (ns)** | -6 | -6 | -6 | -6 |
| **Noise level σ** | 0.0062 | 0.0096 | 0.0079 | 0.0038 |
| **Modulation depth λ** | 0.049 | 0.165 | 0.180 | 0.090 |
| **DEER SNR** | 7.9 | 17.2 | 22.8 | 23.4 |
| **Background Decay η** | 0.09 | 0.32 | 0.35 | 0.31 |
| **Background parameters (a[$ns^{-1}$],b)** | 0.0454 | 0.0662 | 0.0655 | 0.0301 |
| **RMSE (Background subtraction)** | -0.0705<br>$-3.1 \cdot 10^{-5}$ | -0.1935<br>$-1.1 \cdot 10^{-4}$ | -0.2172<br>$-9.8 \cdot 10^{-5}$ | -0.1136<br>$-7.6 \cdot 10^{-5}$ |
| **Spin concentration measured by cw-ESR (µM)** | 32 | 57 | 30 | 18 |

**Table S6: Euclidean geometry analysis of the distance found in a trimer-of-dimers geometry** with the fixed distance AB = BC = AC = 5.3 nm[4] and $A_1A_2 = B_1B_2 = C_1C_2 = 4.1$ nm (see scheme below). The series of distances are calculated with equal angle values of $\alpha = \beta = \gamma$.

| **α=β=γ (°)** | **A1B1=B1C1=A1C1 (nm)** | **A1C2=A2B1=B2C1 (nm)** | **A1B2=A2C1=B1C2 (nm)** | **A2B2=B2C2=A2C2 (nm)** |
|---|---|---|---|---|
| 0 | 1.75 | 5.68 | 5.68 | 8.85 |
| 20 | 2.31 | 4.99 | 6.30 | 8.72 |
| 30 | 2.85 | 4.63 | 6.57 | 8.56 |
| 40 | 3.44 | 4.28 | 6.80 | 8.34 |
| 50 | 4.06 | 3.96 | 7.00 | 8.06 |
| 60 | 4.68 | 3.67 | 7.15 | 7.71 |
| 70 | 5.27 | 3.45 | 7.26 | 7.32 |
| 80 | 5.84 | 3.30 | 7.33 | 6.87 |
| 90 | 6.38 | 3.25 | 7.35 | 6.38 |
| 100 | 6.87 | 3.30 | 7.33 | 5.84 |
| **110** | **7.32** | **3.45** | **7.26** | **5.27** |
| **115** | **7.52** | **3.55** | **7.21** | **4.98** |
| **120** | **7.71** | **3.67** | **7.15** | **4.68** |
| **125** | **7.89** | **3.81** | **7.08** | **4.37** |
| 130 | 8.06 | 3.96 | 7.00 | 4.06 |
| 140 | 8.34 | 4.28 | 6.80 | 3.44 |
| 150 | 8.56 | 4.63 | 6.57 | 2.85 |
| 160 | 8.72 | 4.99 | 6.30 | 2.31 |
| 180 | 8.85 | 5.68 | 5.68 | 1.75 |

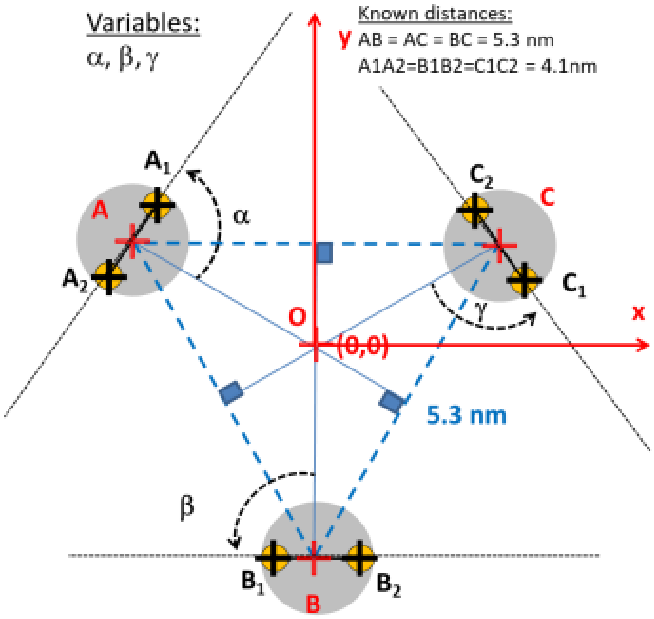

Schematic describing the angle α, β, γ definition used for calculating distances within a TODs array unit. The two-fixed distances in this model are the 4.1 nm separating the PAS domain in a homodimer, and the 5.3 nm distance prescribed by a Tar/Tsr chemosensor array observed by cryo-electron tomography in minicells[4].

**Table S7: Distances measured in the homology model (Figure 5a and 5b) between N5-N5 of Flavin in the PAS domain**. The predicted distance domain $P_{tot}(r)$ presented in Figure 5d is obtained by summing gaussian curves $y_i(r)$ with a center value $r_i$ equal to the distance measured and an amplitude $A_i$ equal to the number of distance measured and a standard deviation σ fixed to 0.15 nm:

$$P_{tot}(r) = \sum_i y_i(r) = \sum_i \frac{A_i}{\sqrt{2\pi\sigma}} \exp\left(-\frac{1}{2}\left(\frac{r-r_i}{\sigma}\right)^2\right)$$

| **Distance measured (Å)** | 31.25 | 33.75 | 36.25 | 38.75 | 41.25 | 43.75 | 51.25 | 53.75 | 56.25 |
|---|---|---|---|---|---|---|---|---|---|
| **Number of distance measured** | 4 | 14 | 8 | 7 | 6 | 9 | 1 | 1 | 4 |
| **Distance measured (Å)** | 58.75 | 61.25 | 63.75 | 66.25 | 68.75 | 71.25 | 73.75 | 76.25 | 78.75 |
| **Number of distance measured** | 2 | 8 | 5 | 13 | 6 | 8 | 5 | 8 | 12 |

**Table S8: 4P-DEER experimental parameters[1] of TarCheWCheA-iLOV *in cell* compared to CheA-iLOV *in vitro* (see Figure 6).** These parameters were obtained after zero-order phasing of the experimental data and background subtraction with the function: $B(t) = exp(a * t^2 + b * t + c)$. The data was then subtracted by this background correction as suggested by Ibáñez et. al.[2]. The time domain was truncated (reported as $t_{max}$). The "Offset Time" (noted as $t_{min}$) was determined using the "correctzerotime.py" function of the Deerlab software[3]. The noise level was evaluated using the standard deviation of the imaginary part of the data divided by the maximum value of the real part. Signal to noise ratio was then deduced by the ratio of the modulation depth λ over the noise level σ. The modulation depth λ was evaluated by subtracting S(t=$t_{min}$) - B(t=$t_{min}$). Finally, the background decay η is obtained by the relation $\eta = 1 - \frac{B(tmax)}{B(tmin)}$.

| | ***In cell*** | | ***In vitro*** |
|---|---|---|---|
| **Parameters :** | **TarCheWCheA-iLOV** | **CheA-iLOV** | **CheA-iLOV** |
| **$t_{max}$ (ns)** | 1486 | 1486 | 1446 |
| **Offset Time $t_{min}$ (ns)** | -2 | -2 | -2 |
| **Noise level σ** | 0.0018 | 0.0015 | 0.0025 |
| **Modulation depth λ** | 0.013 | 0.005 | 0.0044 |
| **DEER SNR** | 4.3 | 1.3 | 0.4 |
| **Background Decay η** | 0.17 | 0.18 | 0.25 |
| **RMSE (Background subtraction)** | 0.0055 | 0.0049 | 0.0042 |
| **Spin concentration measured by cw-ESR (µM)** | 187 | 230 | 209 |

### Supplementary Figures:

#### Figure S1: IPTG-induced expression of Aer measured by *in cell* cw-ESR.

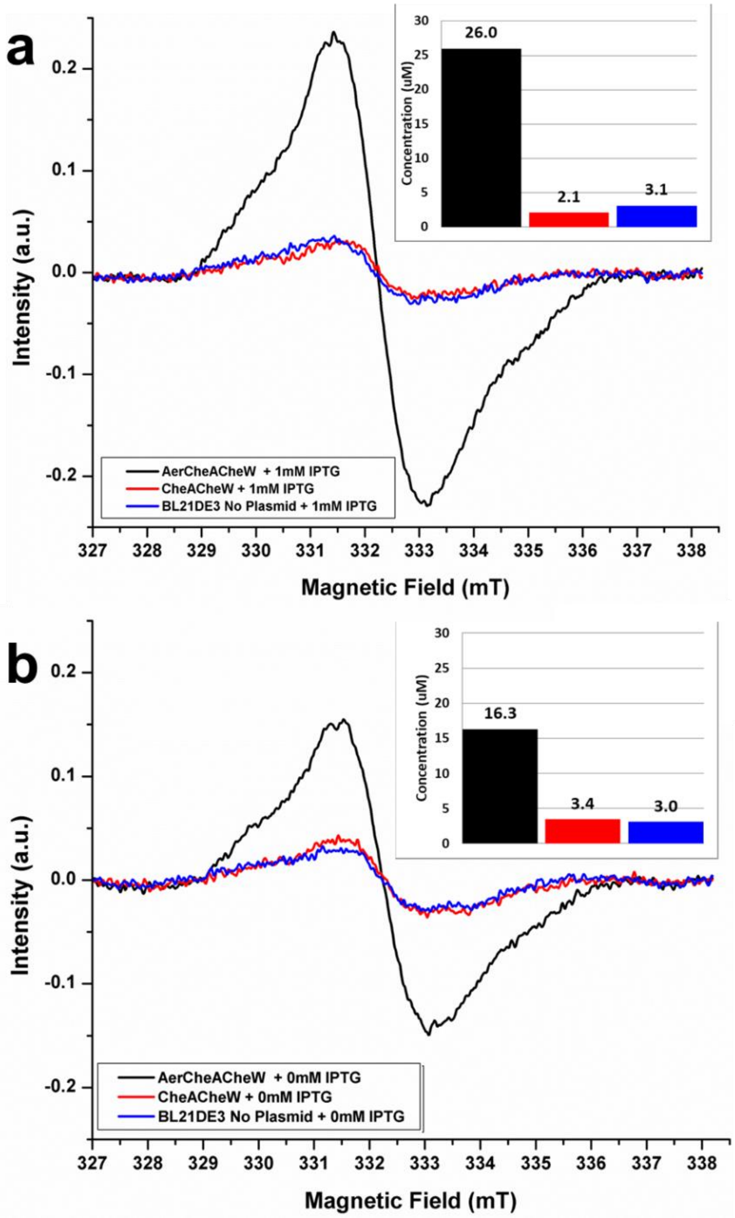

**Figure S1: IPTG-induced expression of Aer measured by *in cell* cw-ESR.** *In cell* cw-ESR spectra of Aer co-expressed with the signaling partner CheA and CheW (black curve), the signaling partners alone (red curve), and *E. coli* BL21-DE3 alone without plasmid (blue curve). Each cell sample was grown then induced with 1 mM IPTG (Isopropyl ß-D-1-thiogalactopyranoside) (panel a) or measured after the same time duration without IPTG induction (panel b). Inset: Column chart of the measured concentration of the flavin semiquinone radical. The concentration was measured by comparing the double integral of the cw-ESR signal against a TEMPO (2,2,6,6,-tetramethyl-piperidin-1-oxyl) calibration scale.

### Figure S2: Cw-ESR spectra for Aer and Aer site-directed variants.

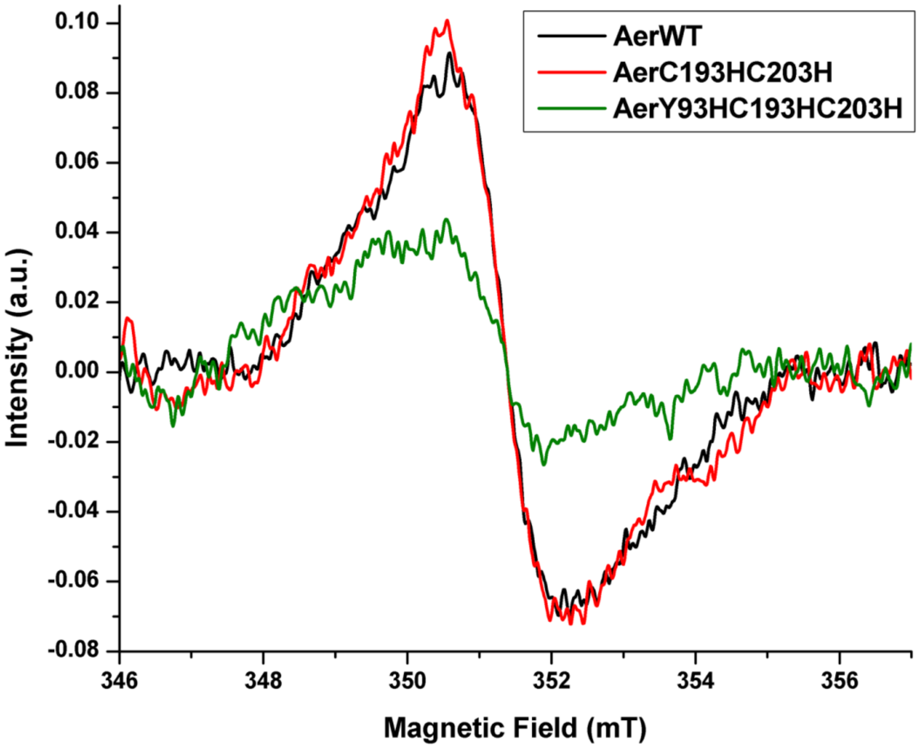

**Figure S2: Cw-ESR spectra of Aer *in cell* without (black curve) or with the residue substitutions C193H/C203H (red curve) and Y93H/C193H/C203H (green curve).** The cysteine substitutions (C193, C203), which are not expected to affect flavin binding, do not change the ESR signal intensity (semiquinone radical), whereas the Tyr93 substitution Y93H produces a large drop in the ESR signal intensity. The Y93H substitution reduces flavin binding[5–7]. It has been shown that overexpression of Aer with IPTG induces the accumulation of flavin in the membrane[5], the cw-ESR spectrum of the Y93H/C193H/C203H variant indicates that free FAD does not appreciably contribute to the ESR signal.

#### Figure S3: *In cell* 4P-DEER analysis of Aer co-expressed with and without CheA/CheW.

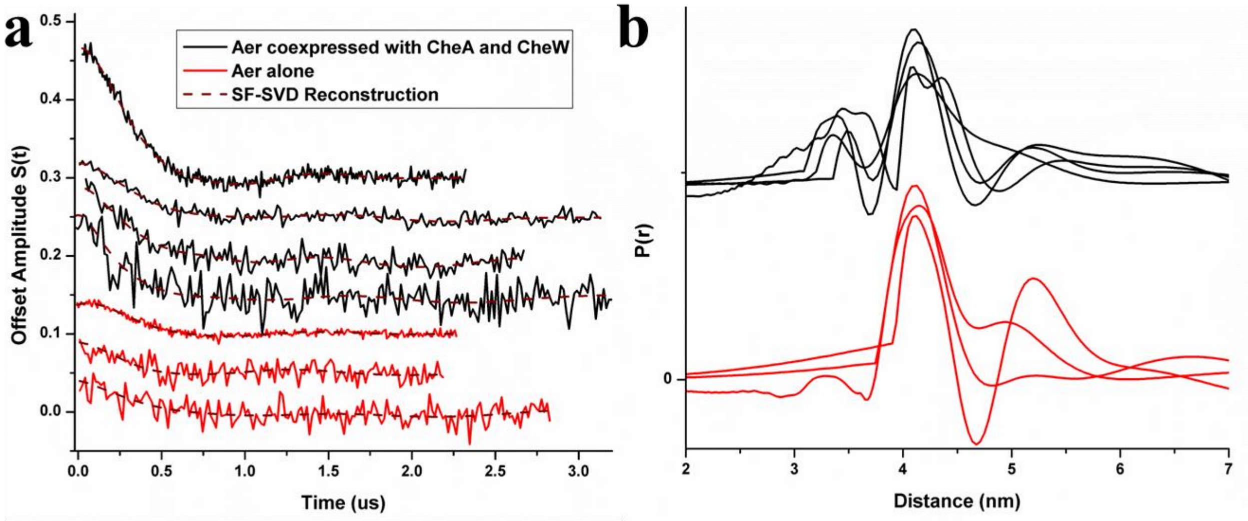

**Figure S3: *In cell* 4P-DEER analysis of Aer in BL21 with coexpression (black) or without coexpression of CheA/CheW (red).** (a) 4P-Deer time domain with different evolution times ($t_{max}$). Modulation depths of $\lambda$ = 0.157, $\lambda$ = 0.049, $\lambda$ = 0.092 and $\lambda$ = 0.092 were measured for Aer($^1H/^{14}N$) with CheA/CheW whereas smaller modulation depths of $\lambda$ = 0.036, $\lambda$ = 0.032 and $\lambda$ = 0.031 were found without CheA/CheW (see Table S2). A greater modulation depth $\lambda$ is an indication of a higher FAD-bound Aer population. (b) 4P-DEER distance domain obtained by using the SF-SVD method[8]. No short distances are observed in the 3-4 nm range for the Aer without coexpression of the signaling partners CheA and CheW.

#### Figure S4: X-band cw-ESR spectra of Aer grown in isotopically enriched media

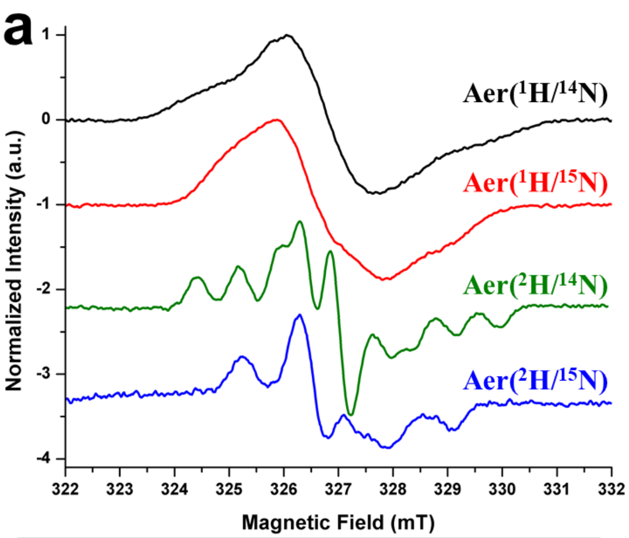

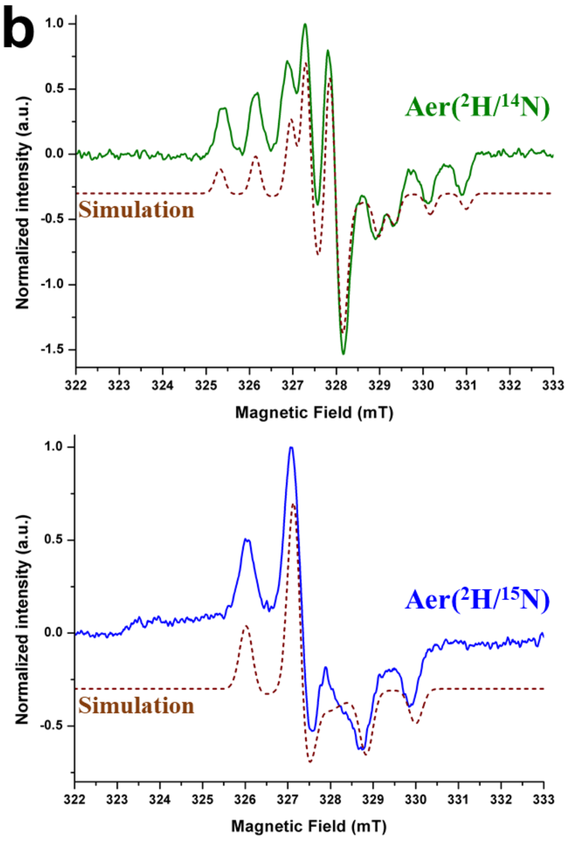

**c**

| | cwESR Global Fitting | | |
|---|---|---|---|
| | $A_{xx}$ (MHz) | $A_{yy}$(MHz) | $A_{zz}$ (MHz) |
| $^{14}$N/($^{15}$N)[(a)] **(Position 5)** | 1.7 (2.4) | 1.2 (1.7) | 56.3 (79.0) |
| $^{14}$N/($^{15}$N)[(a)] **(Position 10)** | 4.5 (6.3) | 4.4 (6.2) | 23.1 (32.4) |

**Figure S4: X-band cw-ESR spectra of Aer grown in isotopically enriched media** (a) X-band cw-ESR spectra of Aer co-expressed with CheA/CheW in BL21 cells grown in different isotopically rich media measured at 293 K; (b) Simulation fits (dash-brown lines) of the experimental cw-ESR spectra of Aer enriched with the isotopes $^2$H/$^{14}$N (green) and $^2$H/$^{15}$N (blue). Simulations were carried out using a global fitting procedure in Easyspin[9] for both $^{14}$N/$^{15}$N isotopes; hyperfine coupling constants (hfcc) were the only parameters fit (see details of the matlab script in Supplementary Information). (c) Values in parentheses are conversion of $^{15}$N hfcc to an equivalent $^{14}$N hfcc via the gyromagnetic ratio $|\gamma_{15N}/\gamma_{14N}|$ = 1.4028.

### Figure S5: Aer($^1$H/$^{14}$N) and Aer($^1$H/$^{15}$N) 3P-ESEEM fitting

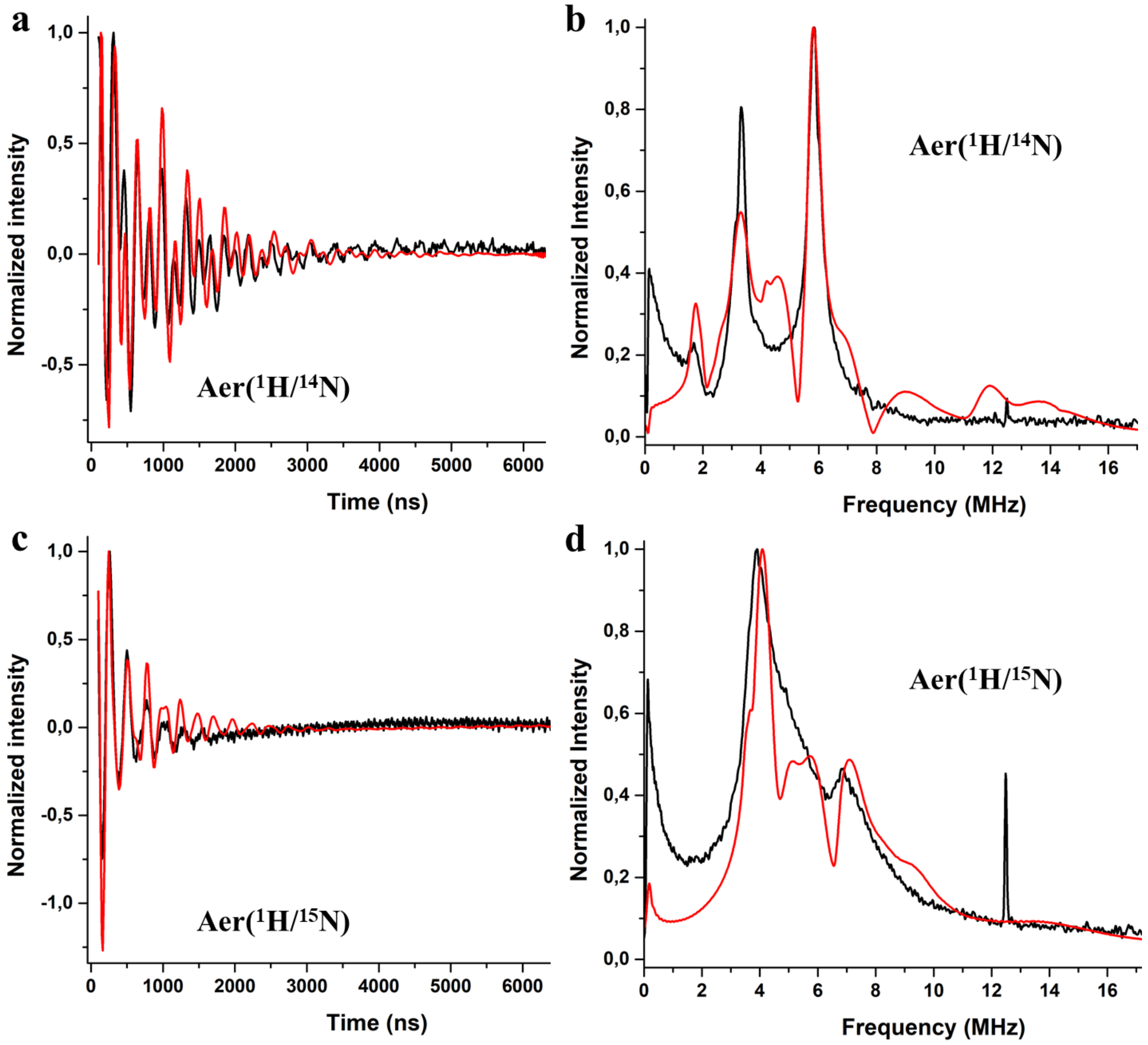

**Figure S5: Aer($^1$H/$^{14}$N) and Aer($^1$H/$^{15}$N) 3P-ESEEM fitting**. Q-Band 3P-ESEEM spectra of Aer($^1$H/$^{14}$N) (a) in the time domain and (c) in the frequency domain. Q-Band 3P-ESEEM spectra of Aer($^1$H/$^{15}$N) (b) in the time domain and (d) in the frequency domain. In each graph, the experimental curves are plotted in black and their respective simulations are in red. The simulation script employed for the fits is given below and the simulation parameters are reported in the **Table S3**. Experimental 3-pulse ESEEM sequence π/2-tau-π/2-T-π/2-tau-echo was employed at a temperature T = 150 K to study the spin modulation of the $^{14}$N and $^{15}$N atoms on the flavoprotein. To avoid any tau artifact, tau was varied from 160 ns to 230 ns by step of 10 ns. For every tau value, the 3P-ESEEM was recorded using a starting mixing time T = 100 ns and a dwell time of 16 ns to have a sufficient frequency resolution in the area of 1-20 MHz. For this pulse experiment, the resonator was critically coupled and a pulse length of 16 ns was employed for typical π/2 pulses. Spectra were simulated with the saffron.m function in EasySpin[9].

#### Figure S6: Temperature profiles of the spin lattice relaxation time $T_1$ of isotopically enriched Aer co-expressed with CheA and CheW *in cell*.

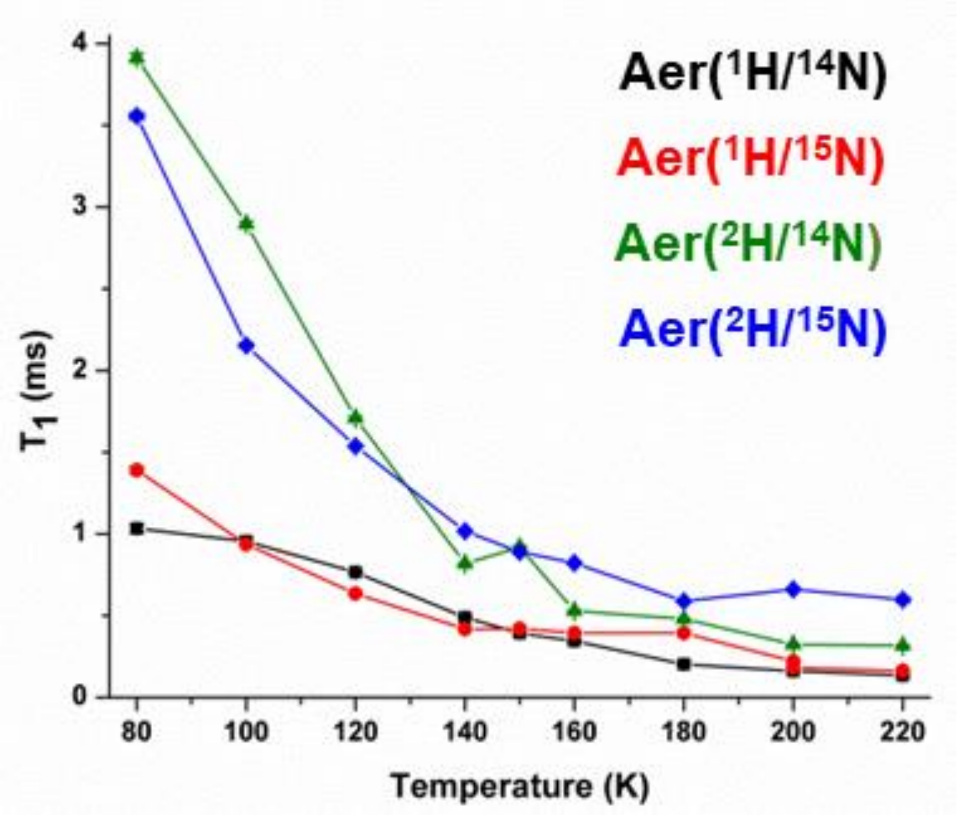

**Figure S6: Temperature profiles of spin lattice relaxation times $T_1$ of isotopically enriched Aer co-expressed with CheA and CheW *in cell*.** $T_1$ for the Aer anionic semiquinone with various enriched isotopes: black square Aer($^1H/^{14}N$), red circle Aer($^1H/^{15}N$), green triangle Aer($^2H/^{14}N$), blue diamond Aer($^2H/^{15}N$). For this pulse experiment, the resonator was critically coupled and typical $\pi/2$ and $\pi$ pulses were 16 ns and 32 ns, respectively.

#### Figure S7: Aer($^2$H/$^{14}$N) and Aer($^2$H/$^{15}$N) Mims $^2$H-ENDOR spectra

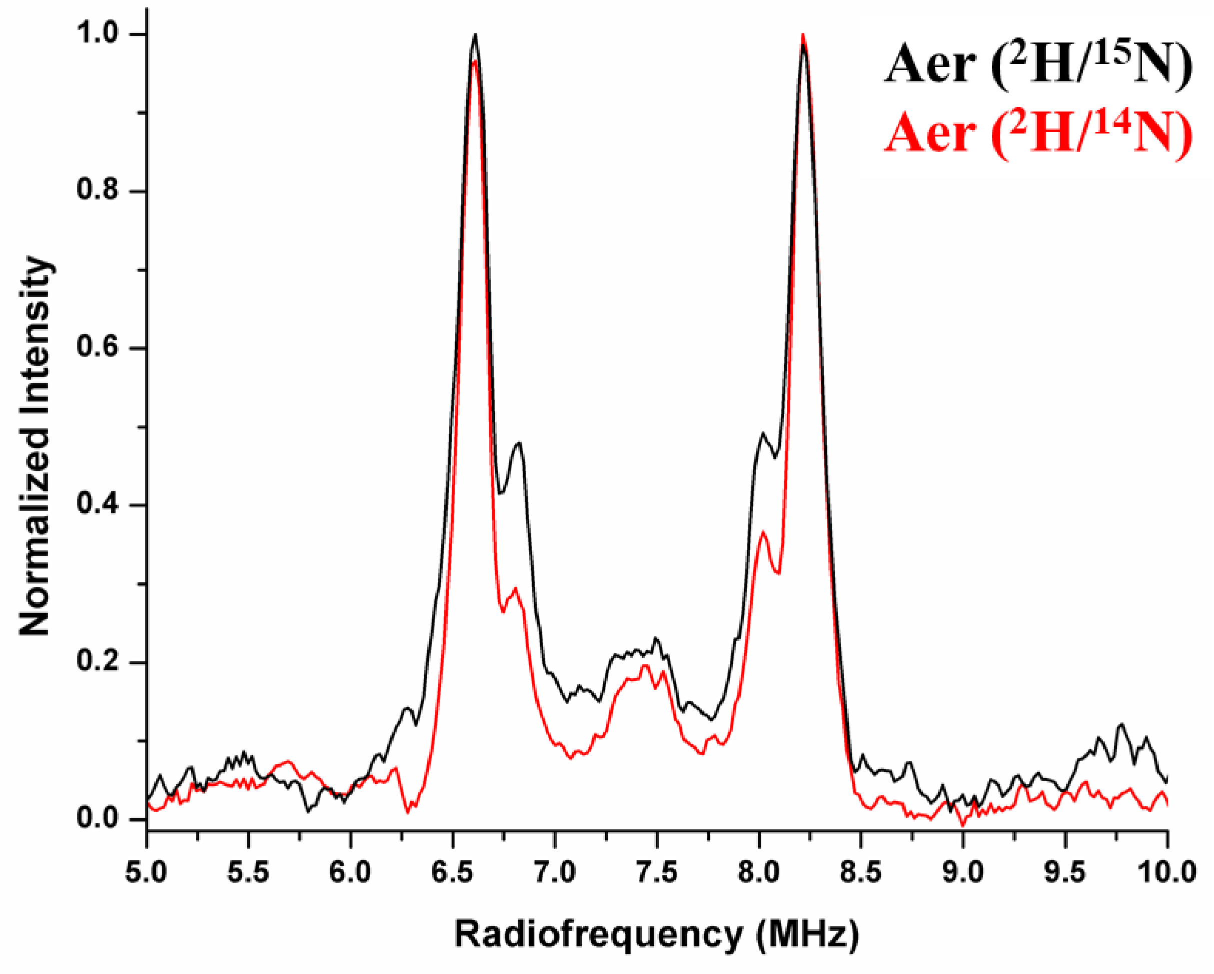

**Figure S7**: **Aer($^2$H/$^{14}$N) and Aer($^2$H/$^{15}$N) Mims $^2$H-ENDOR spectra.** Mims $^2$H-ENDOR of Aer grown in deuterated media co-expressed with CheA and CheW (red curve: $^2$H/$^{14}$N, black curve: $^2$H/$^{15}$N) at a frequency of 33.65102 GHz (the central resonance for $^2$H is expected at around 7.4 MHz). The hyperfine coupling constants **(Table S4)** match the values observed by $^1$H-Davies ENDOR, thereby indicating that deuterium substitution does not alter the flavin electronic structure.

### Figure S8: *In cell* deuterium solvent exchange experiment with Aer($^1$H/$^{14}$N)-CheA-CheW

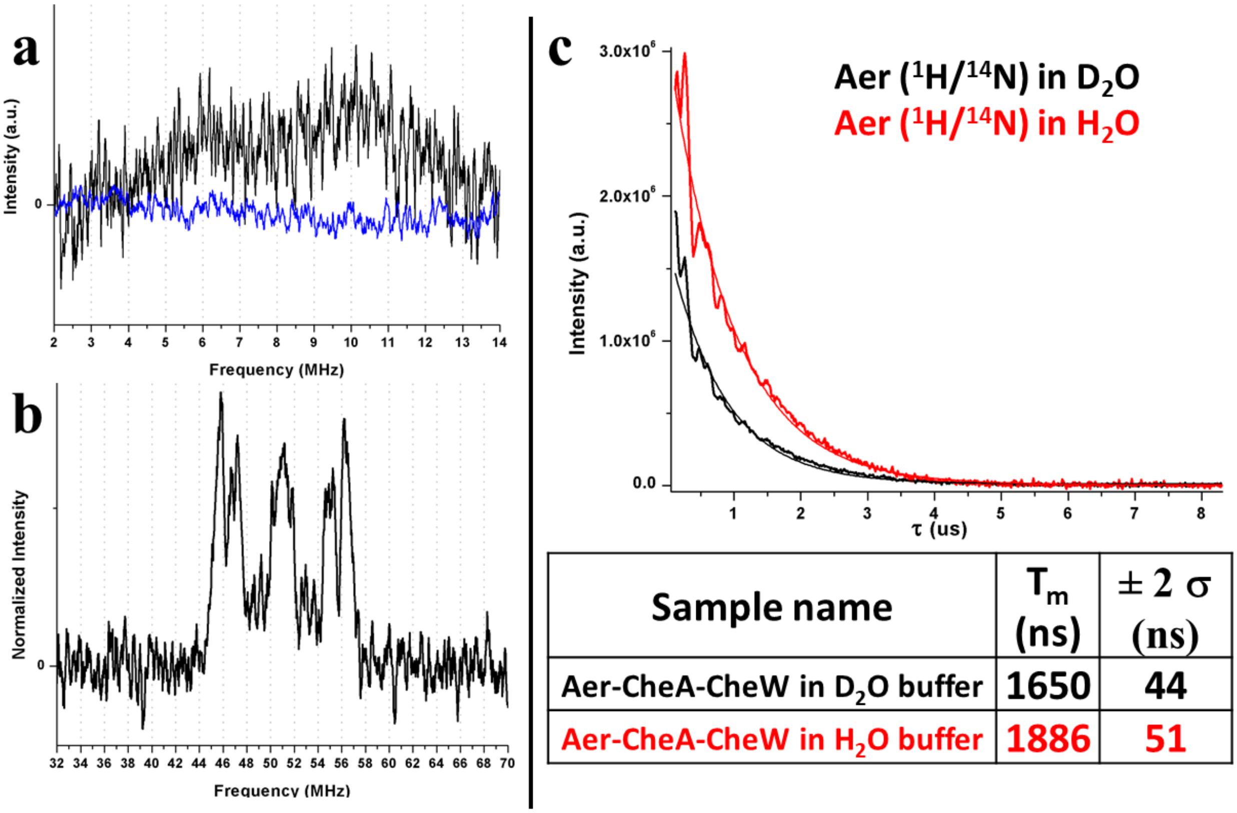

| Sample name | $T_m$ (ns) | ± 2 σ (ns) |
|---|---|---|
| Aer-CheA-CheW in $D_2O$ buffer | 1650 | 44 |
| Aer-CheA-CheW in $H_2O$ buffer | 1886 | 51 |

**Figure S8:** ***In cell*** **deuterium solvent exchange experiment with Aer($^1$H/$^{14}$N)-CheA-CheW.** (a) Aer-CheA-CheW resuspended in $D_2O$/$D_3$-glycerol buffer analyzed by *in cell* $^2$H-ENDOR recorded using the Davies pulse sequence (black curve) or by using the Mims pulse sequence (blue curve). No signal is observed around 7.4 MHz (see **Figure S7** for comparison). (b) Aer-CheA-CheW resuspended in $D_2O$/$D_3$-Glycerol buffer analyzed by *in cell* $^1$H-ENDOR and recorded using the Davies pulse sequence (black curve). (c) Comparison of the relaxation times for the same cell sample containing over-expressed Aer-CheA-CheW dispersed in $H_2O$ buffer (red curve) and in $D_2O$ buffer (black curve). The table summarizes the $T_m$ values obtained by fits to a monoexponential decay function. No clear changes in relaxation time are observed after solvent exchange.

#### Figure S9. Expression and Activity of CheA-iLOV.

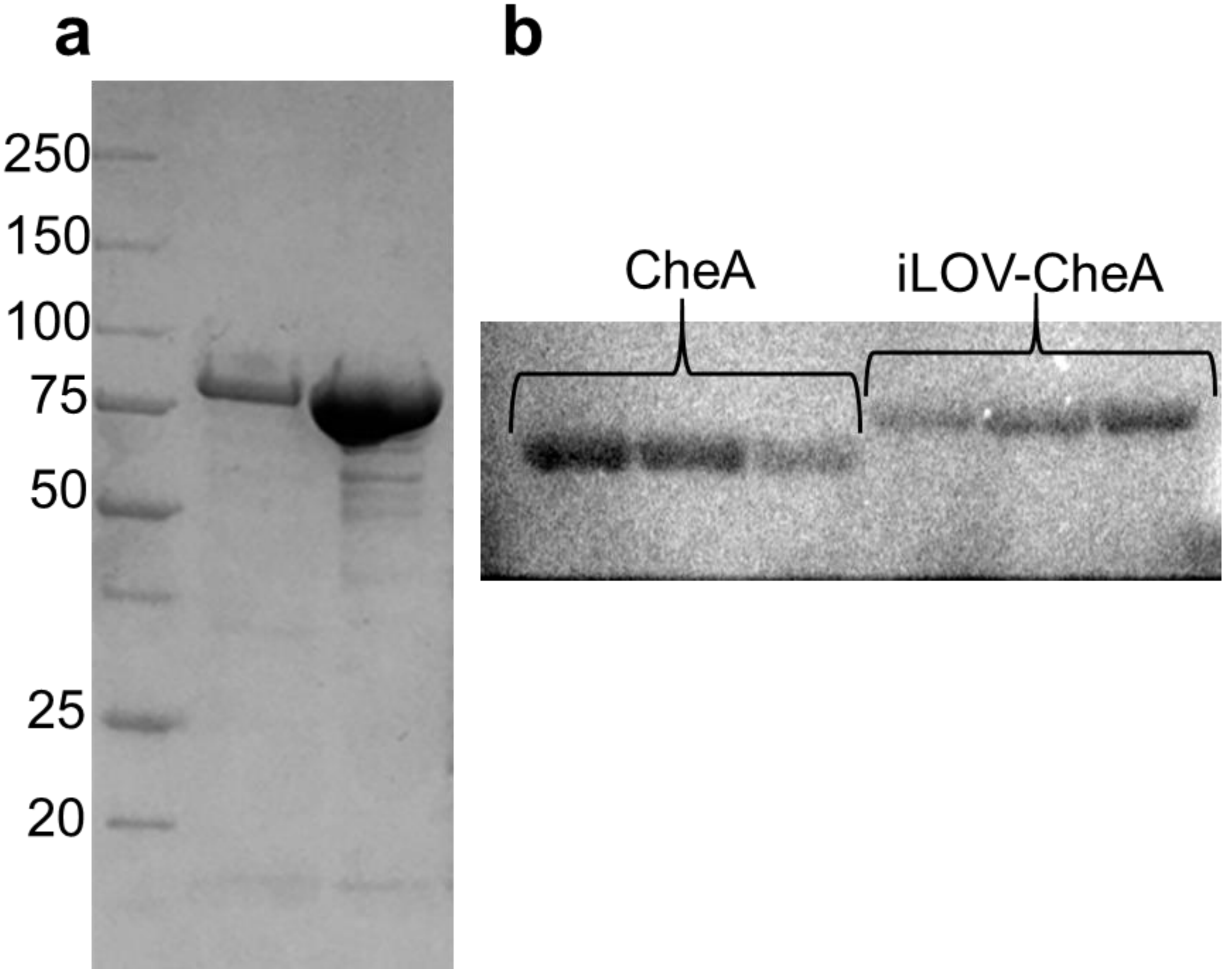

**Figure S9**: **Expression and Activity of CheA-iLOV.** (a) SDS-PAGE gel of CheA-iLOV (molecular weight ~ 87.5 kDa). Left lane is the protein ladder, central lane is the column elution fraction obtained after nickel affinity purification and right lane is the fraction obtained after size-exclusion chromatography. (b) Autoradiogram of CheA autophosphorylation with ATP- $\gamma$ $^{32}$P. The fusion of iLOV to the N-terminus of CheA increases the molecular weight (as evidence by an upshift in the gel) but does not inhibit the autophosphorylation activity of CheA. The experiments were carried out in triplicate, Lane 1-3 is for CheA and Lane 4-6 is for CheA-iLOV.

### Figure S10. Cw-ESR and $^1$H-ENDOR spectra of CheW-CheA-iLOV.

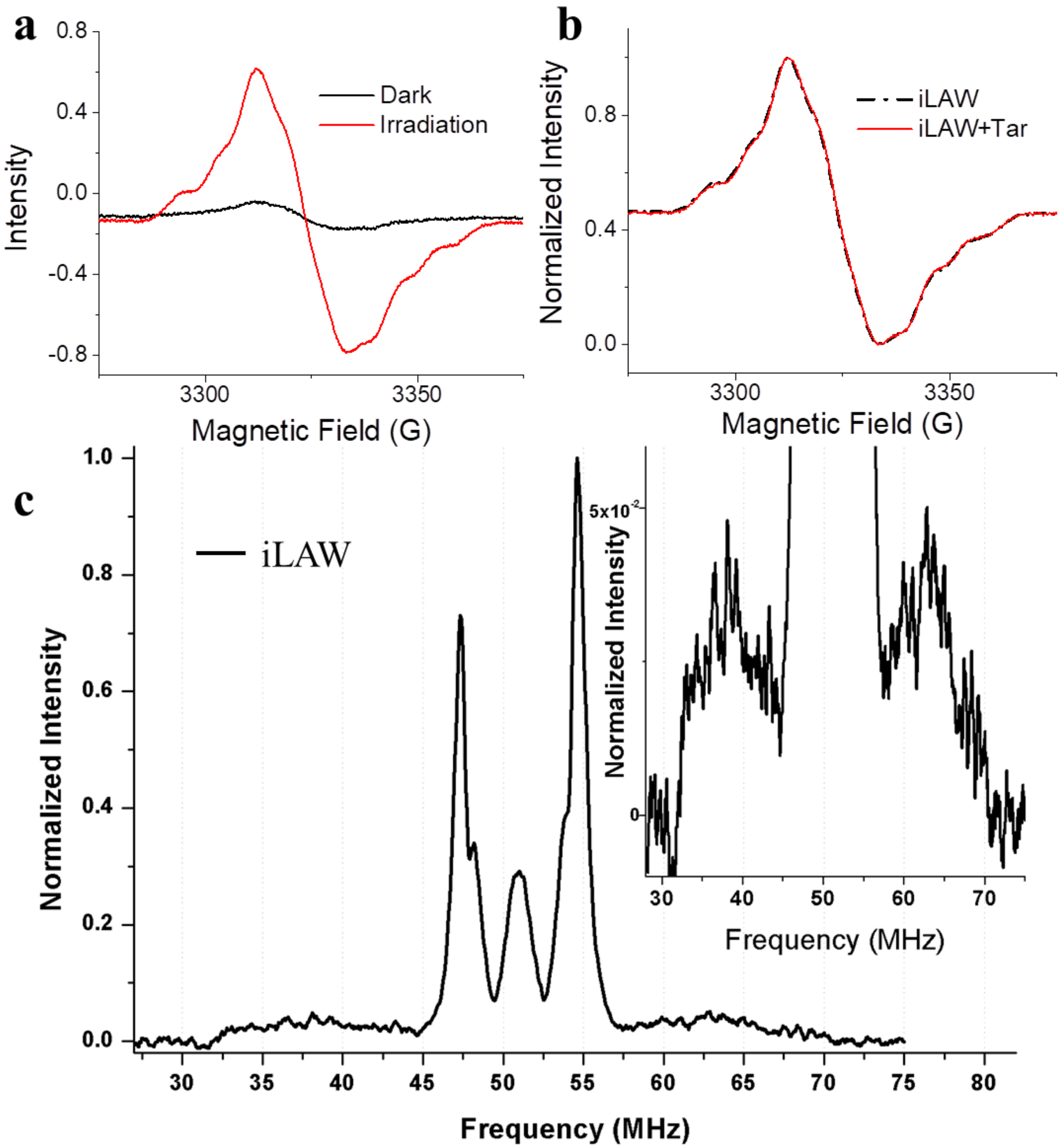

**Figure S10: Cw-ESR and $^1$H-ENDOR spectra of CheW-CheA-iLOV (iLAW).** (a) Cw-ESR spectra of CheA-iLOV co-expressed with CheW in the dark and after blue light irradiation (3 min). (b) Co-expression of CheW-CheA-iLOV along with the Tar receptor does not affect the spectral features of the flavin radical in iLOV (c) *In cell* Q-Band $^1$H Davies ENDOR spectrum of CheW-CheA-iLOV sample at 200 K. Inset: Zoom on the broad component indicates a 25-30 MHz hfcc owing to the $^1$H proton located on the N5 position of the isoalloxazine ring of the FAD neutral semiquinone radical.

#### Figure S11: Kinetics of CheA-iLOV degradation in the cellular environment.

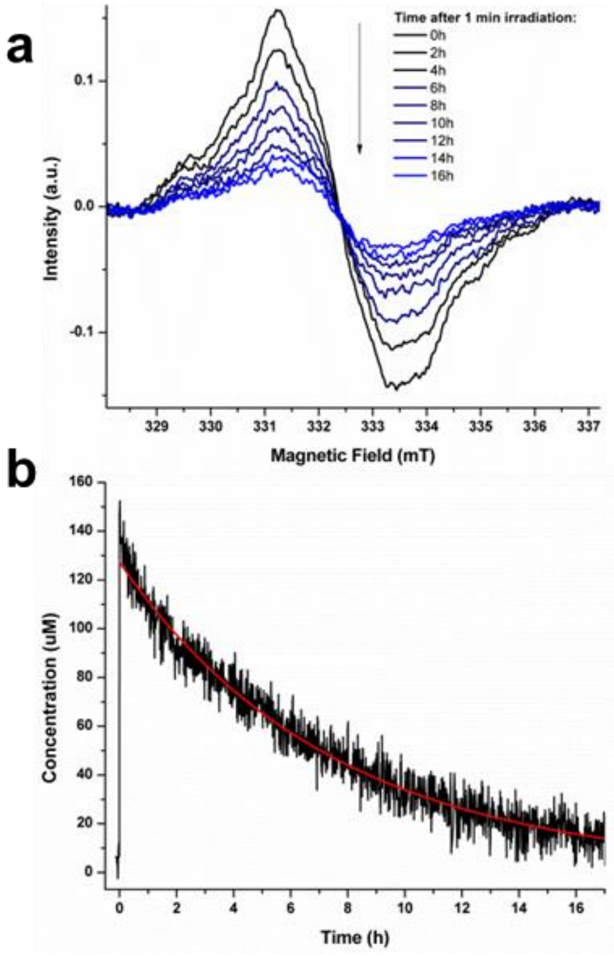

**Figure S11: Kinetic of degradation of CheA-iLOV in cellular environment**. (a) *In cell* cw-ESR spectra of CheA-iLOV measured every 2h at 293 K after 1 min irradiation with blue light ($\lambda$ = 457 ± 10 nm) (b) Spin concentration versus the time after irradiation (black curve). The time was set to zero at the beginning of the irradiation. Cw-ESR spectrum was recorded every 45s. The concentration was measured by comparing the double integral of the cw-ESR signal against a TEMPO (2,2,6,6,-tetramethyl-piperidin-1-oxyl) calibration scale. A monoexponential decay fit (red curve), $y(t) = A_1 * exp(-\frac{t}{t_{decay}}) + A_0$ was obtained with the following parameters: $A_1$ = 126 ± 2 μM, $A_0$ = 1.3 ± 2 μM and $t_{decay}$ = 7.4 ± 0.4 h.

#### Figure S12: *In cell* 4P-DEER measurements of TarCheWCheA-iLOV in different growth conditions in minimal media.

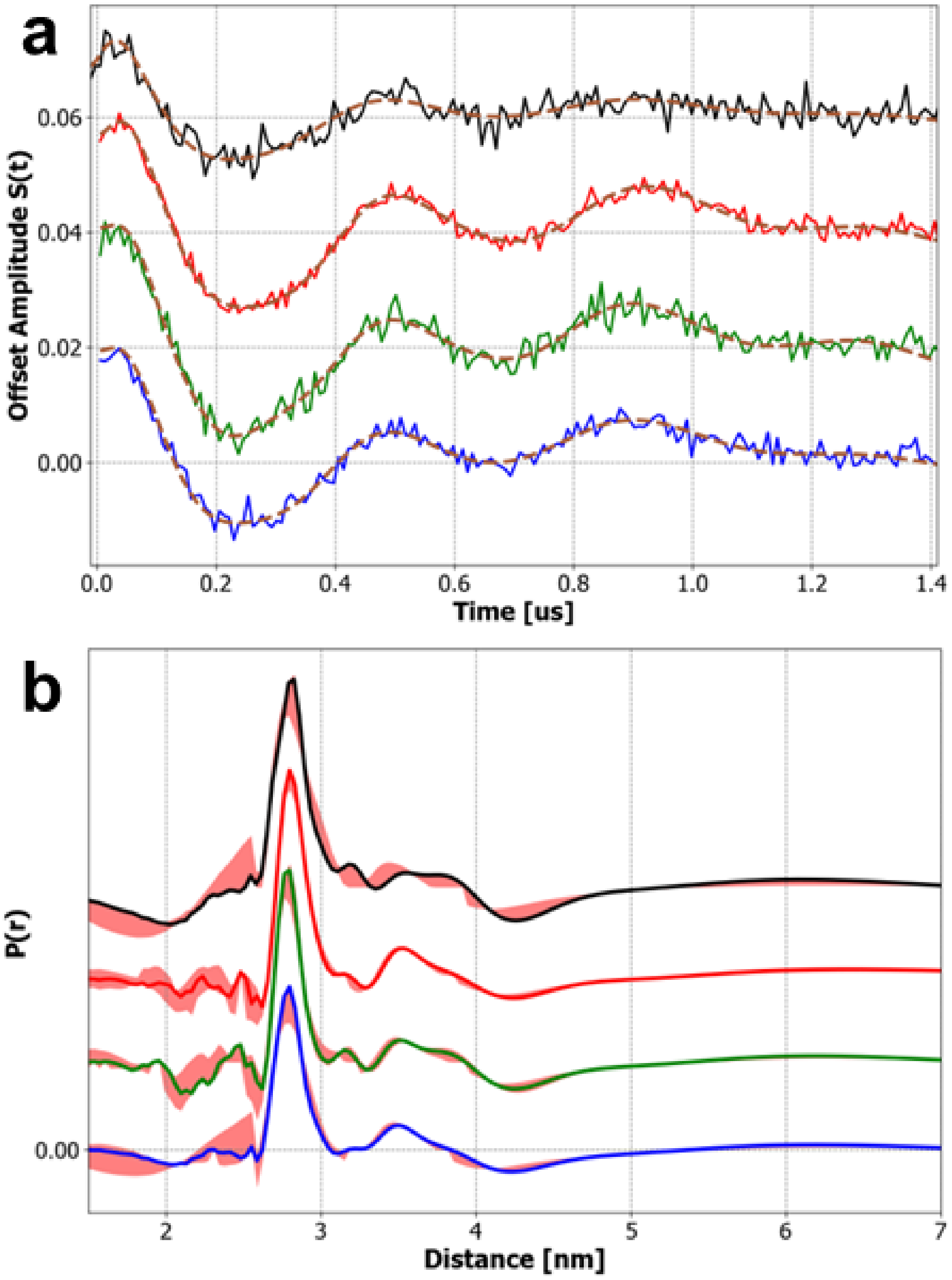

**Figure S12: *In cell* 4P-DEER measurements of TarCheWCheA-iLOV in different growth conditions:** TarCheWCheA-iLOV in minimal media alone (first black curve) with 1mM ATP (second red curve), with 2 mM $Ni^{2+}$ (third green curve) and with 2 mM aspartate (fourth blue curve) (a) Time domain and reconstructed time domain (SF-SVD method[8]) (dash brow line) measured at T = 180 K by Q-Band 4P-DEER sequence and (b) its associated domain distribution P(r) (SF-SVD method). Red shading represents the error in the distance reconstruction and was calculated as described in Srivastava et al.[8].

### MATLAB code for global fitting

1) EasySpin[9] Script used for global fitting simulation of cw-ESR data (**Figure S3**).

```
%%%
%This script is used for globally fitting isotopically modified (14N/15N) cw-ESR %spectrum.
%The parameters employed are fitted via a variable 'Param' inserted in the %classically defined
%spin system 'Sys' and used in a 'user-defined' function (here %named 'Global14N15N_M4G').
%
%Timothee Chauvire November, 11th, 2024
%%%
clear all;
Filename1= 'A040619_AerD2O_M=4G.dat'; %load experimental spectrum Aer(2H/14N)
Filename2 = 'A041019_AerN15D2O_M=4G.dat'; %load experimental spectrum Aer(2H/15N)
temp1 = importdata(Filename1);
temp2 = importdata(Filename2);

B1 = temp1(:,1)/10; % Convert Magnetic Field in mT
spc14N = temp1(:,2)/max(temp1(:,2)); % Normalize the spectra to simulate
B2 = temp2(:,1)/10; % Convert Magnetic Field in mT
spc15N = temp2(:,2)/max(temp2(:,2)); % Normalize the spectra to simulate

% Definition of the 2 set of spin systems to simulate the 14N and 15N hyperfine interaction
Sys14N = struct('g',[2.00436 2.00402 2.00228],'S',0.5,'lwpp',[0.22 0]);
Sys15N = struct('g',[2.00436 2.00402 2.00228],'S',0.5,'lwpp',[0.3 0]);
Sys14N.Nucs = '14N, 14N';
Sys14N.n = [1 1];
Sys15N.Nucs = '15N, 15N';
Sys15N.n = [1 1];
Sys14N.A = [1.666300205       1.217980273    56.34843717;    4.517784059    4.44814188
                   23.12313496]; % define the hfccs of the two 14N used for the simulation

% Adjust the different spin system value by using a set of coefficient parameters
Param = [1.0 1.0 1.0 1.0 1.0 1.0 1.0 1.0 0.84];
Sys14N.Param = Param; % Create a Variable that will be incorporate in the spin system structure
used by EasySpin for the simulation
gyr14N15N = 1.4027; % Gyromagnetic ration for the conversion between 14N and 15N
Sys15N.A = Sys14N.A.*gyr14N15N; % copy the hfccs of the two nitrogens
Sys15N.Param = Param;
% define the experimental parameters:
Exp14N = struct('mwFreq',9.1965,'Harmonic',1,'Temperature',295,...
     'Range',[B1(1) B1(end)],'ModAmp',0.4,'nPoints',1024);
Exp15N = struct('mwFreq',9.1925,'Harmonic',1,'Temperature',295,...
     'Range',[B2(1) B2(end)],'ModAmp',0.4,'nPoints',1024);
Exp = Exp14N;

% define the optimization parameters used by the function pepper
Opt.Verbosity = 2;
Opt.nKnots = 20;
Opt.Method = 'hybrid';
Vary.Param = [0.4 0.4 0.4 0.4 0.4 0.4 0.2 0.2 0.2];
spc = [spc14N;spc15N] ; % concatenate the 2 experimental spectrum Aer(2H/14N) and Aer (2H/15N) in
one variable 'spc'
Exp.nPoints = 2048; % Use the size of the variable 'spc' for the fit

% Execute the fitting procedure by employing an homemade function 'Global14N15N_M4G'
esfit('Global14N15N_M4G',spc,Sys14N,Vary,Exp,Opt)

%%%
%This function is used as a global fit procedure for simulating isotopically
%modified (14N/15N) cw-ESR spectrum.
%The parameters employed are stored in a variable called 'Param'.
%The trick is then to duplicate the 14N spin system to a 15N spin system and adjust %the fit via
%the variable 'Param'.
%The two spectra simulated are then concatenated via an output variable y.
%
```

```
%Timothee Chauvire November, 11th, 2024
%%%
function y = Global14N15N_M4G(Sys,Exp,Opt)
fullSys14N = Sys; %extract the spin system used for the fitting
fullSys15N = fullSys14N; %duplication of the spin system for Aer(2H/15N) spectrum
Param = Sys.Param; %extract the fitting parameters
%Adjust the hfccs tensor and linewidth of the simulated spectrum with the fitting parameters
fullSys14N.A(1,1) = Sys.A(1,1)*Param(1);
fullSys14N.A(1,2) = Sys.A(1,2)*Param(2);
fullSys14N.A(1,3) = Sys.A(1,3)*Param(3);
fullSys14N.A(2,1) = Sys.A(2,1)*Param(4);
fullSys14N.A(2,2) = Sys.A(2,2)*Param(5);
fullSys14N.A(2,3) = Sys.A(2,3)*Param(6);
fullSys14N.lwpp = Sys.lwpp*Param(7);
Exp.nPoints = 1024;
gyr14N15N = 1.4027; % Gyromagnetic ration for the conversion between 14N and 15N

fullSys15N.A = fullSys14N.A.*gyr14N15N;
fullSys15N.Nucs = '15N,15N';
fullSys15N.lwpp = Sys.lwpp*Param(8);

[x14N,y14N] = pepper(fullSys14N,Exp,Opt);
Exp.mwFreq = 9.166; %Compensate the magnetic field shift via a frequency shift
[x15N,y15N] = pepper(fullSys15N,Exp,Opt);
y = [y14N'; Param(9).*y15N']; %Use of a parameter for a relative scaling of the spectra
end
```

2) EasySpin scripts used for a fit of the 3P-ESEEM data (**Figure S4**).

```
%%%
%This script is used for fitting 3PEseem spectrum in frequency domain (or %alternatively in time
%domain with a Background subtraction step).
%Two homemade functions are used for the fit: 'FrequencyFitEseem' or %'TimeFitEseem'.
%
%Timothee Chauvire November, 11th, 2024
%%%
clear all
Filename1 = 'A051019_N15Aer.dat';
Filename2 = 'A051019_N15Aer_Time.dat';
freq = importdata(Filename1);
time = importdata(Filename2);

tau = 0.21;
Exp.Sequence = '3pESEEM';
Exp.Field = 1192.8;
Exp.dt = 0.016;
Exp.tau = tau;
Exp.T = 0.1;
Exp.mwFreq = 33.67;
Exp.ExciteWidth =5e8;

Sys.g = [2.00436 2.00402 2.00228];
Sys.Nucs = '15N,15N';
Sys.A = [3.4      0.1 84.9;  3.3 2.9 36.7];
Sys.T1T2 = [4000,200];

Opt.nKnots = 480; % A lot of orientation are needed as the hfccs of the 14N/15N are strongly
anisotropic
Opt.TimeDomain = 0 ;
Opt.ProductRule = 0;
Opt.ZeroFillFactor = 4;
Opt.Method = 'matrix';
Opt.Verbosity = 2;
Vary.A = [2 2 20;4 4 20];

Exp.nPoints = size(freq(:,2),1)/2; % A fit on half of the frequency domain was achieved
spcFreq = freq(1:length(freq)/2,2)/max(freq(1:length(freq)/2,2));
```

```
esfit('FrequencyFitEseem',spcFreq,Sys,Vary,Exp,Opt);

%Either a fit in frequency or a fit in the time domain can be used:
%Exp.nPoints = size(time(:,2),1)/2;
%spcTime = time(1:length(time)/2,2)/max(time(1:length(time)/2,2)) ;
%esfit('TimeFitEseem',spcTime,Sys,Vary,Exp,Opt);
________________________________________________________________________________________

%%%
%This function is used as a global fit procedure for simulating 3PEseem spectrum in
%frequency domain.
%
%Timothee Chauvire November, 07th, 2019
%%%
function fout = FrequencyFitEseem(Sys,Exp,Opt)
[x,y,out] = saffron(Sys,Exp,Opt); %Simulate the 3P-ESEEM spectrum in the time domain but extract the frequency domain via the 'out' variable
% Get the limit of half of the positive quadrant in the frequency domain
N1 = length(out.fd)/2 ;
N2 = length(out.fd)/4 ;
fout = abs(out.fd(:,N1+1:N1+N2))/max(abs(out.fd(:,N1+1:N1+N2))); % extract the data in the frequency domain and normalized it
end
________________________________________________________________________________________
%%%
%This function is used as a global fit procedure for simulating 3PEseem spectrum in
%time domain with a background subtraction (Exponential of a polynomial function).
%An Homemade function called 'bcgd_eseem' is used for the background subtraction.
%
%Timothee Chauvire November, 07th, 2019
%%%
function y_new = TimeFitEseem(Sys,Exp,Opt)
Exp.nPoints = 512;
[x,y,out] = saffron(Sys,Exp,Opt);
xnew = 1000.*x; % Convert the ns time scale in us
[y2,background]= bcgd_eseem(xnew,real(y),3); % Homemade function to subtract an exponential polynome in the time domain
y_new = y2/max(y2);
end

%%%
%This function enables to fit the background via an exponential of a nth degree
%polynomial.
%Three inputs are needed : x (the x absciss), y (the y absciss), and nPoly the
%order of polynomial needed for the fit.
%Two outputs are generated : the new values of y with background subtraction y_new,
%and the 'background' variable used for the subtraction.
%
%Timothee Chauvire November, 07th, 2019
%%%
function [y_new,background]= bcgd_eseem(x,y,nPoly)
ylog = log(y); % convert the time domain in logarithmic scale
p = polyfit(x,ylog,nPoly); %polynomial fit in the logarithmic domain
f = polyval(p,x); %reconstruct the polynomial function in the logarithmic domain
y_new = exp(ylog)./exp(f)-1; % subtract the background
background = exp(f); % export the background
end
```
